# Diverse perceptual biases emerge from Hebbian plasticity in a recurrent neural network model

**DOI:** 10.1101/2024.05.30.596641

**Authors:** Francesca Schönsberg, Davide Giana, Yukti Chopra, Mathew Diamond, Sebastian Goldt

**Author notes:** M.D. and S.G. contributed equally.

## Abstract

Perceptual biases offer a glimpse into how the brain processes sensory stimuli. While psychophysics has uncovered systematic biases such as contraction (stored information shifts towards a central tendency), and repulsion (the current percept shifts away from recent percepts), a unifying neural network model for how such seemingly distinct biases emerge from learning is lacking. Here, we show that both contractive and repulsive biases emerge from continuous Hebbian plasticity in a single recurrent neural network. We test the model in four different datasets, two sensory modalities and three experimental paradigms: two working memory tasks, a reference memory task, and a novel “one-back task” that we designed to test the robustness of the model. We find excellent agreement between model predictions and experimental data without fine-tuning the model to any particular paradigm. These results show that apparently contradictory perceptual biases can in fact emerge from a simple local learning rule in a single recurrent region of the brain.

## Introduction

The seemingly simple task of categorising the intensity of a stimulus is prone to several perceptual biases. For example, in a tactile intensity working memory task, where the subject is presented with two sequential stimuli on each trial and is required to report whether the second stimulus was weaker or stronger than the first (fig. 1a), both human and rodent subjects display a *contraction bias*: they tend to overestimate the strength of the first stimulus if it lies below the weighted average of past stimuli, and underestimate the first stimulus if it lies above the average [1–4]. A similar contraction occurs in tasks where subjects are asked to report the size or intensity of a stimulus after some time has elapsed since exposure to the stimulus [5, 6]. By contrast, in intensity reference memory tasks [7–10], where the subject reports whether one stimulus per trial was perceived as strong or weak, a *repulsive bias* emerges: subjects are more likely to characterise a stimulus as strong if the previous trial’s stimulus was weak, and vice versa [10].

**Figure 1:**
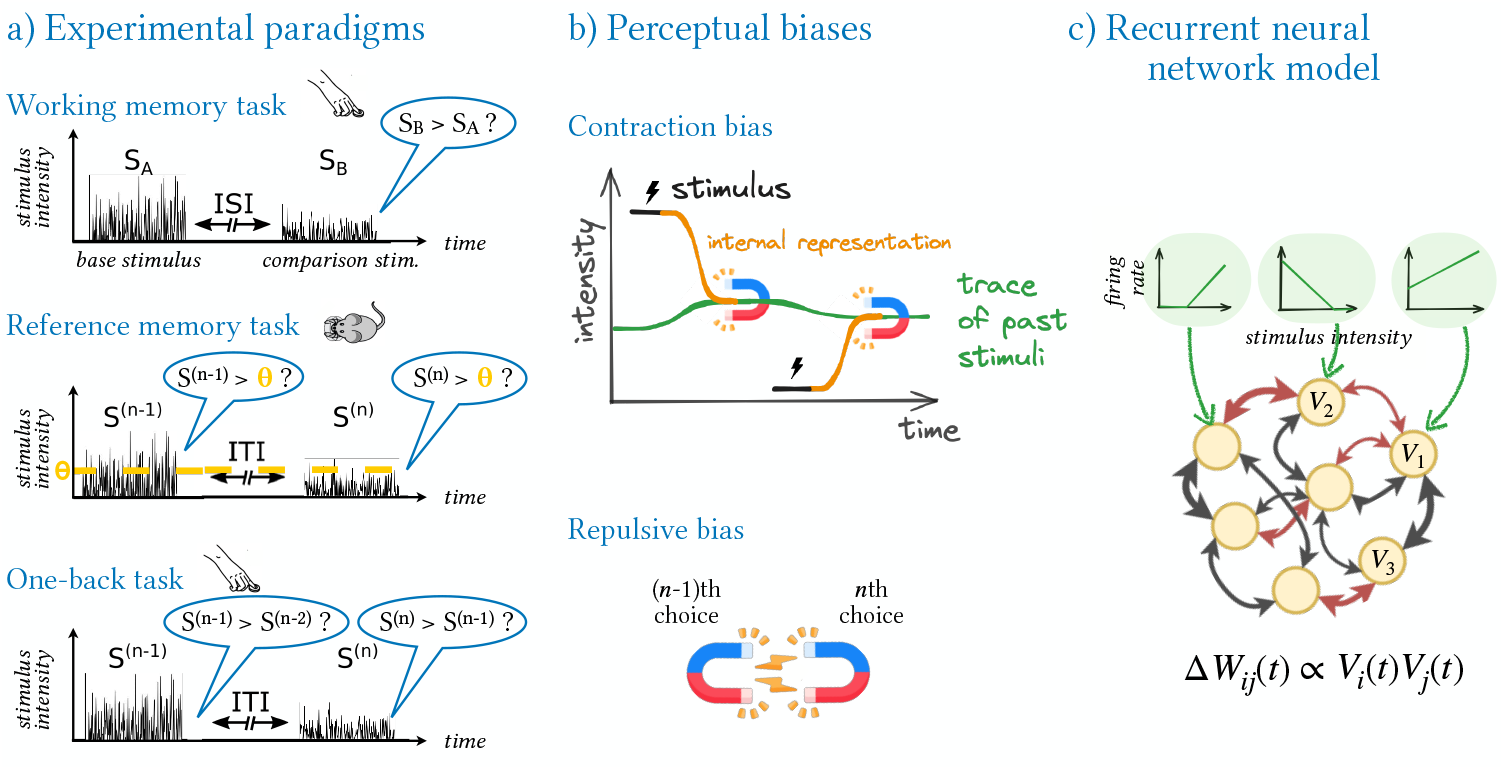
A single neural network model explains two perceptual biases in three perceptual memory paradigms. **a)** We consider **three experimental paradigms**: a *working memory* task in humans, where subjects are tasked with reporting whether the second of a pair of sequential stimuli of strengths *S*_*A*_, *S*_*B*_, separated by a variable inter stimulus interval (ISI), was stronger than the first; a *reference memory* task in rats, where animals compare the strength of one stimulus per trial to a fixed category boundary (orange); and a novel *“one-back” task* that we design to test the generality of our model. Here, human subjects are required to compare the strength of one stimulus per trial to the strength of the stimulus in the preceding trial. In all three paradigms, trials are separated by a variable inter-trial interval (ITI) and subjects receive rewards after correct choices. **b)** The performance of humans and rodents in these paradigms reveals **two perceptual biases**: the *contraction* of stimulus representations to the trace of past stimuli, and the *repulsion* between successive judgments. **c)** We reproduce both perceptual biases in **a single neural network model**, where a fully-connected recurrent neural network (RNN) is driven by external inputs (green) encoding stimulus intensity, while its recurrent connections are continuously reshaped by Hebbian plasticity and can be positive or negative (red/black).

These biases are of interest because they offer a window into the computational processes underlying perceptual judgements. On the experimental side, a great number of psychophysical studies, comprising different sensory modalities, species, and details of experimental design, have provided a detailed characterisation of both contraction and repulsion [3, 5, 7–12]. On the modelling side, comprehensive phenomenological models have been built to account for these biases [1, 10, 13–17]. Nevertheless, the neural mechanisms at work remain poorly understood. While the idea that neural circuits could perform working memory tasks through a quasi-continuous line of fixed points dates back to the 1990^*′*^s [18, 19], Boboeva *et al*. [20] showed only recently that a neural network model can reproduce the specific perceptual biases observed by Akrami *et al*. [3]. However, existing theoretical models have static ad-hoc connectivity and are tuned to reproduce a single specific bias. Further, they overlook learning – the boundary must be learned in reference memory categorisation, as evidenced by the notable performance disparities of trained subjects between the onset and the end of experimental sessions [10]; current models have failed to capture how perceptual biases arise in parallel with short-timescale learning. Finally, while recent phenomenological models point to the existence of general principles underlying diverse perceptual biases [17], there exists no single neural network model that, without reconfiguration, can account for multiple biases expressed across different experimental paradigms.

Here, we show that a single recurrent neural network (RNN) with ongoing Hebbian plasticity learns stimulus representations that reflect the perceptual biases of humans and rodents. We first show how the model reproduces contraction and repulsive biases, as well as the history-dependence of choices, as measured in a novel tactile working memory experiment that we perform with human subjects. In addition, we show that our model also reproduces the biases observed in two previously published data sets, the auditory working memory task of Akrami *et al*. [3], and the tactile reference memory task of Hachen *et al*. [10]. Our analysis shows how both perceptual biases emerge from a plastic attractor driven by recurrent dynamics and Hebbian plasticity. We finally design a new experimental paradigm that tests the generality of the model. In this “one-back task”, fig. 1a, participants have to compare the strength of the stimulus presented on each trial to the stimulus of the previous trial, making each stimulus serve both as comparison and, successively, as base. We find a novel, bi-modal contraction bias in the performance of human subjects and reproduce it in the RNN model. We stress that we do not fine-tune the model to any individual paradigm, nor do we fit the RNN to experimental data or use gradient descent for learning. Instead, we fix hyper-parameters across experiments and use only local Hebbian plasticity to let the weights evolve continuously.

## Results

As an overview, we first illustrate the key experimental paradigms, the perceptual biases that emerge in such paradigms, and the schematic architecture of the neural network model (fig. 1).

### A recurrent neural network model with ongoing Hebbian plasticity

#### Network architecture

We consider a single recurrent neural network (RNN) model, taking inspiration from neurobiological evidence indicating that perceptual biases may occur in a recurrent subregion of the motor cortex (see *Methods* for details). The network is composed of *N* threshold linear units, which model the neurons or the ensemble activity of a group of them. The activity state of the network 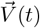 represents the firing rate of all *N* units at each time *t*. Although the network state evolves continuously through the synchronous update of all units, for clarity we present the update for each unit *V*_*i*_ individually, which obeys the equation [21–23]:

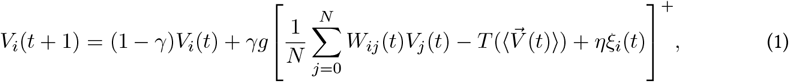

where *γ* sets the timescale of the neuronal dynamics and [*z*]^+^ = max(0, *z*). The neural input field, within the square brackets, is based on Treves [24] and Schönsberg *et al*. [25] and determines the activity of the neurons through three components: the inputs from the other neurons weighed by the recurrent connectivity *W*_*ij*_(*t*) (which represents the strength of the synapse from neuron *j* to neuron *i*), a time-dependent threshold 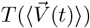, and an external input *ξ*_*i*_(*t*), *d*escribed below. The threshold does not represent any specific biophysical quantity directly; instead, it serves to maintain a tendency towards a desired average activity value, preventing activities from diverging towards zero or infinity. Its value depends on the average activity across all units, 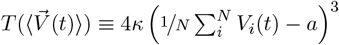 as in [25], where a and *ϰ* are parameters setting the desired average and the contribution of the threshold respectively (see *Methods* for their values), while the cubic function is a simple supra linear function that preserves the sign of the deviation.

#### Synaptic plasticity

For simplicity, we assume that the network is dense, with bidirectional connections *W*_*ij*_(*t*) between all pairs of neurons, defined in a connectivity matrix 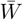. All the connections evolve in parallel to the neural dynamics. Specifically, starting from random initial weights, the weights follow the Hebbian plasticity rule [26], here reported again for one single connection:

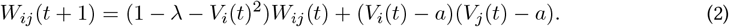

The parameters *λ* and *V*_*i*_(*t*)^2^ are simple normalization terms that ensure the stable operation of a basic Hebbian learning rule, with the latter inspired by Oja’s normalization [27]. The activity values in the second term, instead, are thresholded to yield both positive and negative weights. The key point is that the weights follow a biologically plausible learning rule based on Hebb’s principle of synapse reinforcement, rather than being optimised using a gradient-descent based machine learning algorithm [28]. The key distinctions between these two approaches lie in locality, continuity, and supervision. The Hebbian learning rule is both unsupervised and local, with changes in synaptic strength occurring solely due to the relationship of activity at pre- and post-synaptic neurons. These changes are gradual and continuous, driven by the network’s ongoing activity. In contrast, the optimization-based approaches that our model avoids rely on substantial amounts of information to train weights globally, requiring supervision and multiple iterations. While the Hebbian dynamics of eq. (2) may change the sign of some weights in an apparent violation of Dale’s principle, we observed that this occurs infrequently (see appendix B for details). We also note that recent experimental observations have opened the possibility of a switch in the excitatory/inhibitory effect of a neuron on its postsynaptic target [29].

We fix the hyperparameters *γ, λ, η, g, ϰ, a* to values that put the network in a regime where the recurrent dynamics and the learning occur without divergences (see *Methods* for details). We stress that the model will be tested against the experimental results of all four datasets with the same set of parameters.

#### Stimulating the RNN

At each time step of the neural dynamics of eq. (1), the external stimulus delivered to the units 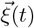 can either be 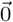 (if no stimulus is applied) or a pattern that represents the intensity of the vibrational or auditory stimulus from the experiment. We designed a set of input patterns 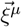 for representing stimulus intensities between *S*^MIN^ and *S*^MAX^ where the index *μ* = 1,…, P runs over the different discrete values of the stimulus intensity. In a scheme that reflects key coding properties of the rat’s primary vibrissal somatosensory cortex (vS1) [30, 31], the input 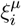 to the *i*th RNN unit for a stimulus of strength *S*^*μ*^ is a threshold linear function of the stimulus strength with a random positive or negative slope, as shown in fig. 1c (green inputs) and *Methods*. This choice reflects the presence of neurons in vS1 whose tuning curves are either positively correlated or anti-correlated with stimulus strength. These conditions are similar to the binary morphing patterns of Blumenfeld *et al*. [32], in that our inputs 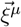, even though they are continuous rather than binary, also display gradual morphing: the correlation between two patterns 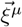 and 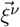 decays with the absolute difference in the corresponding stimulus intensities *ΔS =* |*S*^*μ*^ − *S*^*ν*^ |. This morphing structure will be key in shaping the dynamics of the RNN, as we discuss below. The precise stimulation protocols, i.e. the duration of each stimulation and each inter-stimulus interval for all the tasks are given in appendix A.

#### Reading out the intensity encoded by the RNN and detecting the choice

While the readout of the stimulus intensity represented by the RNN is independent of the task, the way we model the choice depends necessarily on the nature of the experiment. Inasmuch as the decision making derives in humans by verbal instruction and in rats by an extensive training regime, it is reasonable to assume that learning of the behavioral context cannot be captured by model parameter in the current implementation. Specifically, we read out **the intensity encoded by the RNN** state 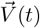 by computing the normalised cosine similarity between 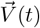 and all the P input patterns 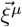. This “overlap profile” has *P* values, which, due to the correlation structure of the input patterns, follow a smooth, bell shaped distribution with a single maximum. The stimulus intensity corresponding to the pattern 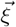 with the maximal overlap is referred to as the network’s internal representation. The **choice** is instead determined based on the experimental task. If the task requires comparing the current stimulus to a previous stimulus, as in the working memory and one-back experiments, we extract the choice at the second stimulus onset by comparing the second stimulus to the network’s internal representation, which will reflect the previous stimulus. If the task instead requires categorising the intensity of a single stimulus, as in the reference memory experiments, the choice is extracted by considering how the internal representation evolves after the stimulus is removed. If the internal representation evolves towards higher stimulus intensities, the stimulus is considered weak, and vice versa.

### Human participants and the RNN model display contraction bias in a working memory task

We first applied the model to a human working memory experiment (fig. 1a). Participants were presented with pairs of vibratory stimuli of strengths (*S*_*A*_, *S*_*B*_) and tasked with reporting which of the two stimuli was stronger, earning a reward for correct choices (see *Methods* for details). This paradigm has been used in several forms to explore working memory [2, 3, 19].

We conducted the experiment with sixteen human subjects, each completing approximately 1000 trials. The performance of human participants (fig. 2a) shows intriguing asymmetries. For example, the pair (105, 80) yielded 21 percent lower performance (69 vs 90 percent) than the pair (80, 105), even if the difference between stimuli is of the same magnitude (25 mm/s). This asymmetry can be interpreted as an outcome of *contraction bias* [3, 5] as follows: during the ISI the memory of the intensity of the base stimulus *S*_*A*_ moves towards a sort of average of all previous stimuli, weighted by their recency through an unknown function. Thus, for the first pair, the memory of *S*_*A*_, ideally fixed at 105, moves closer to the stimulus it must be compared to, *S*_*B*_ (80), making the task harder and leading to errors. For the second pair, the memory of *S*_*A*_ moves toward the recency-weighted average, which is itself in the neighbourhood of 80 mm/sec. Because the base stimulus is in the vicinity of the recency-weighted average, little contraction occurs, leading to fewer errors. Indeed, the *S*_*A*_ memory may be “stabilized.” The thin black arrows in fig. 2a point towards the stimulus pair that is easier to discriminate among pairs with the same difference in stimulus strength. This bias is also manifest in the average psychometric curves across subjects after separating trials according to whether *S*_*A*_ = 80mm/s or *S*_*B*_ = 80mm/s (fig. 2b). Unbiased, perfect choices would result in a step function, which is approached by the psychometric curve obtained when the first stimulus *S*_*A*_ = 80mm/s. If instead the second stimulus is equal to the mean stimulus, *S*_*B*_ = 80mm/s, performance is poor.

**Figure 2:**
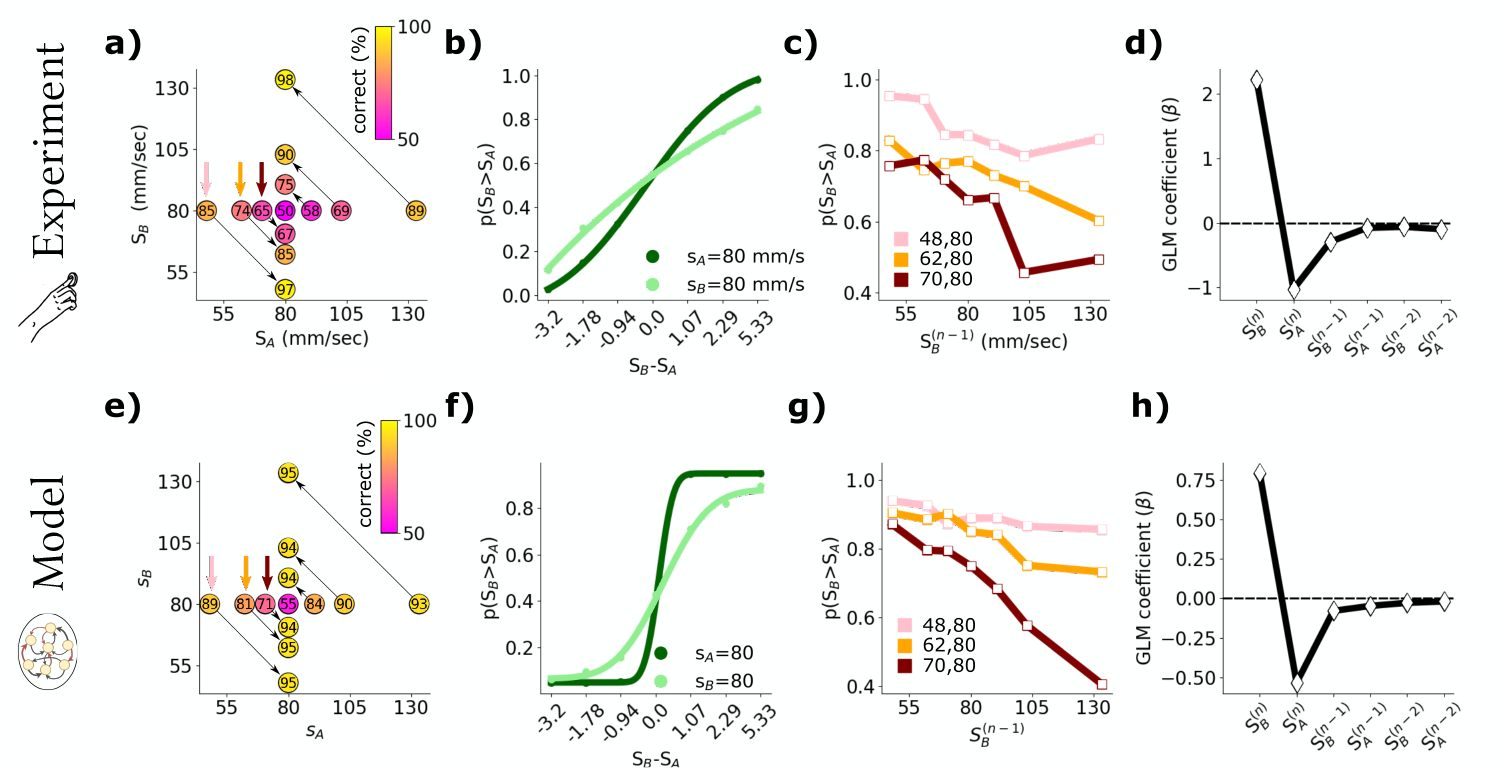
Contraction bias and history-dependence of choices in a working memory task. **a)** Accuracy of human participants in comparing the intensities of two vibrations (*S*_*A*_, *S*_*B*_). Arrows explained in main text. **b)** Psychometric curves based on the stimulus difference if either the first or the second stimulus is 80 mm/sec (light and dark green, respectively). Points correspond to experimental data, solid lines to a sigmoidal fit (see Methods). **c)** Probability that the second stimulus is (correctly) classified as stronger as a function of the strength of the second stimulus *S*_*B*_ in the preceding trial. This plot is shown for the three stimulus pairs highlighted by coloured arrows in a). **d)** Coefficient values (*β*) of a GLM fit of the choice at the *n*th trial as a function of the most recent six perceived stimuli. **e–h)** The same results are reproduced in the RNN model under conditions mimicking those of **a–d)**.

To simulate an experiment, we start both recurrent and Hebbian dynamics, eqs. (1) and (2), from random initial conditions. We then deliver a sequence of stimulus pairs with the same distribution of intensities and ISI’s as in the experiment with human subjects (see *Methods* for details). After 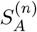 –the base stimulus in trial *n*– has been delivered, the external input 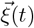 is set to 0 and 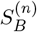 –the comparison stimulus in the same trial– is delivered after a delay. If at the onset of 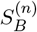, its intensity is larger than the internal representation of the first stimulus, as encoded by the RNN, the RNN makes the choice 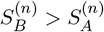, and vice versa. Collecting choices in this simple way, we find clear evidence of contraction bias in the accuracy of the choices of the RNN (fig. 2e), revealing that the internal representation of the RNN contracts towards the average of past stimuli. As seen for human participants, the model’s psychometric curves show good accuracy of the RNN on trials where the first stimulus intensity is near the mean stimulus intensity, while accuracy is lower on trials where the second stimulus intensity is near the distribution mean (fig. 2f). The quantitative differences in the shape of the curves between fig. 2b and f may stem from several factors, including the number of subjects. However, our main focus is to highlight the qualitatively similar behaviour, specifically the steeper increase in response around the mid-stimulus in one condition (dark green) compared to the other (light green).

### History-dependence of choices in working memory task

A fundamental feature of contraction bias reported in previous studies [3, 33] is recency – the dependence of choices on the stimuli of the immediately preceding trial. We uncover this effect in the current data set in fig. 2c. The coloured lines show the probability of making the choice 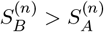 for the three different pairs in the nth trial as a function of the second stimulus in the (*n* − 1)th trial, 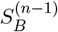. For all three pairs of stimuli shown in fig. 2c, the weaker the preceding stimulus 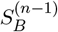, the higher the probability that 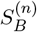 was correctly classified as stronger than 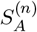. This shows that the stimulus 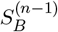 is strongly weighted in the continuously updating average towards which 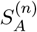 is contracted. Tracing the influence of the six most recently received stimuli, i.e. 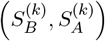 for *k = n, n* − 1, and *n* − 2, on the choice at trial *n* using the coefficient values *β* of a generalised linear model (GLM) reveals a positive effect of the last stimulus (meaning that subjects correctly used the current trial’s *S*_*B*_ intensity to make choices) but a negative effect of previous stimuli; this effect decays for stimuli that are further back in time (see fig. 2d). This is replicated in the model. As shown in fig. 2g, choices made based on RNN representations have the same history-dependence on previous choices as do the choices of human subjects, which is further confirmed by a GLM analysis of RNN based choices (fig. 2h).

In appendix C, we also reproduce the experimental results from a published dataset by Akrami *et al*. [3], which corresponds to a different working memory experiment involving a distinct sensory modality, an auditory delayed comparison task. Specifically, we reproduce the contraction bias and history dependence of choice observed in that experimental design, as well as the relationship between contraction bias and the time interval between the two stimuli.

### Recurrent dynamics and continuous plasticity yield perceptual biases

How do the perceptual biases arise in the RNN model? AAs shown in the supplementary videos and captured in the snap shots in fig. 3a)–d), at the beginning of a session (fig. 3a), when the connectivity is still close to its random initialisation, the RNN develops a large overlap with its first external input (violet line) as seen by the overlap profiles representing the cosine similarity of a given network state with all the different input patterns (going from light grey to dark grey as time increases). However, as soon as the external input is released, the RNN quickly decorrelates and the overlap profile becomes almost flat (fig. 3b). In other words, no particular stimulus intensity is represented by the network. After a few more trials, the effect of Hebbian plasticity becomes apparent: overlap profiles remain peaked even after the stimulus offset, meaning that the network represents a uniquely identifiable stimulus intensity every time, the one corresponding to the peak of the black bell-shaped curve in fig. 3d. This behaviour reflects the emergence of an attractor state with a peaked overlap profile. For *binary* morphing patterns and one-shot learning of the connectivity, Blumenfeld *et al*. [32] indeed proved that a unique attractor emerges. In our case, the attractor emerges from the interplay of continuous Hebbian plasticity, the morphing input pattern structure and intermittent stimuli presented, making it harder to analyse. However, we can make the attractor explicit at each time step *t* by freezing the connections 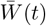 and letting the network converge to the fixed point 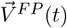 of the recurrent dynamics (eq. (1)). The overlap profiles of the attractors at the sample frozen steps are shown in green in fig. 3a-d (light to dark as time increases); see also supplementary videos.

**Figure 3:**
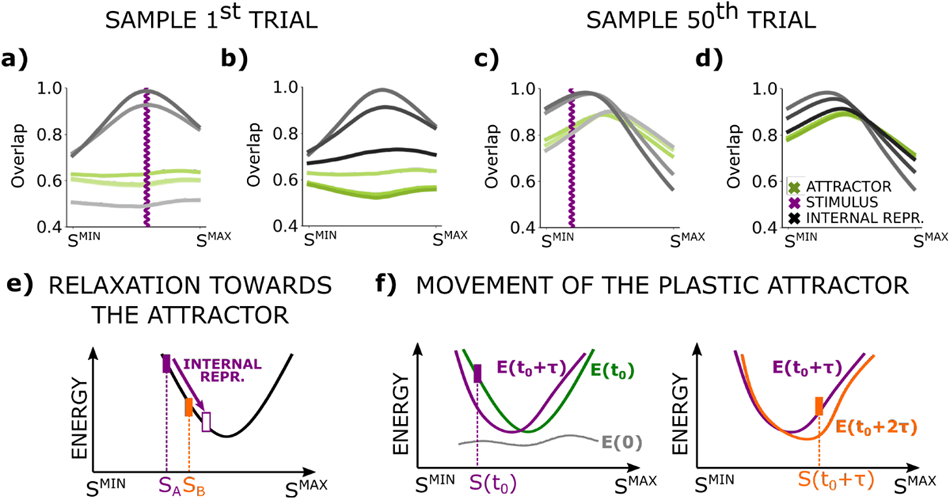
Dynamics of a neural network model that captures perceptual biases. **Top row:** Dynamics of the RNN during the 1st and the 50th trials visualised via the “overlap profile”, i.e. the normalised cosine similarity of the RNN state 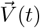 with the input patterns 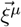 corresponding to each input intensity *S*^*µ*^. We show six different snapshots of the overlap taken at different times (grey lines, increasingly dark as time increases) during and after stimulation (specifically, we use the Reference Memory task introduced later and take the values at the (10,14,20) & (20,36,49) time steps of each trial). The green lines (increasingly dark as time increases) represent the overlap profile of the attractor or fixed point of the RNN dynamics, eq. (1), with the weights fixed at 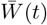. The intensity of the external stimulus is indicated in violet. Refer to videos *V*1 and *V*2 for a visualization of the dynamics of the first and 50th trials. **a)** During a given stimulus input intensity, the RNN state increases its overlap with that stimulus. **b)** After the stimulus is released, the overlap profile quickly decays to an almost uniform profile. **c) & d)** At the 50th trial, the continuous Hebbian plasticity has reshaped the connectivity, leading to the emergence of an attractor with peaked overlap profile. **Bottom row:** Contraction arises from relaxation towards a dynamic attractor, which is reshaped by continuous plasticity. We sketch the energy landscape of the RNN near its minimum as a function of a given set of network states 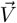 representing different stimulus strengths. **e)** After stimulus *S*_*A*_ is delivered (filled violet bar), the dynamics of the network let the internal representation of that stimulus relax towards the bottom of the energy landscape, which corresponds to the attractor, whose intensity is the trace of past stimuli. In a working memory task, a second stimulus *S*_*B*_ (orange) that is stronger than *S*_*A*_ can be misclassified as weaker if, by the time of its delivery, the memory representation of the first stimulus (empty violet) has descended along landscape beyond *S*_*B*_. **f)** Sketched energy profiles of the attractor at various times: At *t* = 0, the energy *E*(0) is not biased towards any specific stimulus representation (grey). As stimuli are delivered, the energy *E*(*t*_0_) starts having a clear minimum at a specific intensity. Each further stimulation continuously shifts the position of the attractor, i.e. the minimum of the energy, via Hebbian plasticity. Note that the phenomena sketched separately in **e)** and **f)** occur simultaneously.

The attractor constrains the dynamics of the network. The movement of the attractor due to the Hebbian plasticity and the incoming stimuli lead to choices that exhibit perceptual biases. The argument can be simplified by first considering a network with fixed weights, at a certain time *t*. The position of the attractor associated with these weights, and thus the stimulus intensity it encodes, can be viewed as the minimum of an effective energy function *E(t*) in the vicinity of its minimum, as sketched in fig. 3e-f. We do not provide an explicit form of the energy function, as is done for simpler models [24, 34]. Instead, our goal is to offer a conceptual sketch that aids understanding, where the attractor is envisioned as the bottom of a valley, with the dynamics of the RNN following the contours of this valley. At the offset of the first stimulus *S*_*A*_ in a working memory trial (violet in fig. 3e), the state of the RNN evolves to minimise the energy function. The stimulus intensity encoded by the RNN relaxes towards that corresponding to the attractor. In the trial sketched in fig. 3e, this relaxation reduces the distance between the internal representation of the first stimulus and the intensity of the second stimulus, *S*_*B*_ (shown in orange). If the inter-stimulus interval is long enough for the internal representation to “overtake” the intensity of the second stimulus, the RNN makes an erroneous choice by classifying the intensity of the second stimulus as lower than the first.

Although the attractor’s location will generally be close to the mean stimulus intensity, ongoing Hebbian plasticity causes its position to be continuously influenced by the external stimuli. We refer to the *trace of past stimuli* as the quantity tracked by the attractor’s location. While it acts as a sort of recency-weighted average of past stimuli, the precise relationship between past stimuli and attractor location is complex and nonlinear, as detailed in the Discussion. fig. 3f illustrates the effect of two stimuli delivered one after the other, the first weaker and the second stronger than the attractor minimum. At the beginning of a session at *t =* 0, the energy *E*(0) is almost flat. Once a sequence of trials has taken place (*t* = *t*_0_), the energy has a definable minimum emerged thorough Hebbian learning. If a stimulus is then delivered (violet) this induces a change in the energy, which we illustrate in the energy profile *E*(*t*_0_ + *τ*) after some time *τ*. The same occurs with consequent stimuli (orange). The movement of the attractor leads to the history-dependence of the choice. In the working memory experiment, if the second stimulus of the (n 1)th trial was strong, the attractor will have shifted towards higher stimulus intensities, so the internal representation of the first stimulus in the next trial will relax towards the new attractor and might “overtake” the second stimulus, *S*_*B*_ = 80, leading to the patterns of misclassification seen in fig. 2g. Recall that the accuracy of the correct choice for (70, 80) was high if 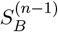 was weak and accuracy declined for progressively stronger values of 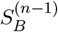. If instead the second stimulus of the (*n* − 1)th trial was weak, the attractor will have shifted towards lower stimulus intensities, facilitating the correct classification of 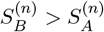.

### Reference memory

In this section, we show that the concurrent dynamical processes in the RNN model – the movement of the attractor and the relaxation of the RNN representation towards it – can also account for the repulsive biases observed in a reference memory task in humans and rats, without any tuning of the hyper-parameters *γ, ϰ, a, λ*. In particular, we show that the serial dependence of choices in *contraction* bias is (perhaps counter-intuitively) an instance of *repulsive* biases.

Experimental data are from the reference memory study of Hachen *et al*. [10], where the rat was tasked with reporting whether each single presented stimulus was weak or strong across a series of trials (fig. 1a). Since Hachen *et al*. [10] showed that in well-trained rats the internal representation of the threshold θ separating weak from strong stimuli was primarily influenced by the distribution of stimuli, rather than the stimulus/reward contingency, we simulate the task by driving the RNN using a stimulus distributions akin to the experimental one, without modelling reward. The model’s choice occurs at the stimulus offset: if the internal representation, i.e. the peak of the overlap profile of the RNN, relaxes towards higher intensities, the delivered stimulus is read out weak, whereas if it relaxes towards lower intensities, the delivered stimulus is read out as strong.

Hachen *et al*. [10] showed that rats internally construct a representation of *θ* to categorise stimuli over successive trials and do not reliably retain it across sessions: the psychometric curves derived from the first three trials of each session deviates only marginally from chance level as compared to those derived from three random trials (fig. 4a). This poor initial performance highlights the importance of learning the network connectivity *dynamically* for any neural network model of this behaviour. Indeed, an initial set of trials is also necessary in our RNN model, to form an attractor that is correlated with past inputs. Consequently, during the first three trials, neuronal configurations relax towards states that are not yet correlated with the effective inputs, resulting in decision probabilities closer to chance level (fig. 4e).

**Figure 4:**
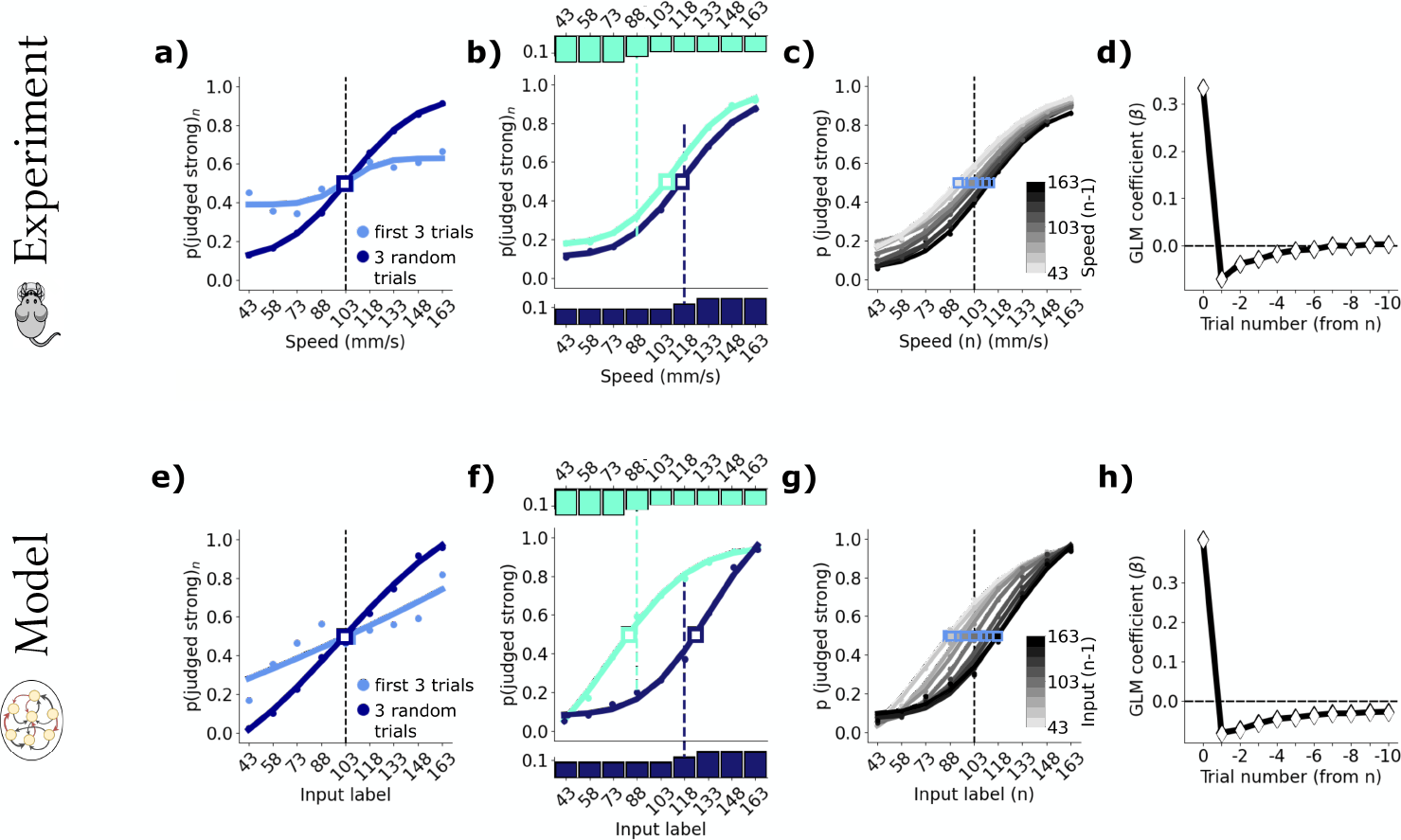
Psychophysical and RNN results in a reference memory task. In **a–d)**, the experimental data from Hachen *et al*. [10] are replotted. **a)** Psychometric curves obtained from the first three trials of each session (light blue) and from three randomly selected trials per session (dark blue). The dashed vertical lines in a-c) mark the category boundary. The point of subjective equality (PSE) is denoted by a square. **b)** Psychometric curves for sessions where input distributions had an average stimulus intensity of 88 (light blue-green histogram) and 118 (dark blue histogram). The input stimuli “attract” the psychometric curves towards the mean of the distribution. This curve shift does not occur if the distribution of the inputs is kept uniform but the reward is delivered according to a category boundary displaced from the mean of the distribution (not reported here, see [10]). **c)** The probability of categorising a stimulus as “strong” conditioned on the intensity of the previous stimulus. Blue squares represent the PSE. **d)** Coefficient values β derived from the generalised linear model (GLM) predicting choice given the stimulus intensities of the ten preceding trials. **e–h)** show results from the RNN under conditions mimicking those of the experiments. In **a–b)** and **e–g)**, points correspond to experimental data, solid lines to sigmoidal fits (see Methods).

The repulsive bias in the reference memory task can be seen in two ways. Figure 4b shows the psychometric curves of rats that were exposed to stimulus distributions biased towards weak and strong stimuli (small light and dark blue histograms, respectively). The psychometric curves in fig. 4b show that an increased frequency of strong stimuli shifts the point of subjective equality (PSE) towards stronger stimuli, and vice-versa for an increased frequency of weak stimuli. The same occurs considering the psychometric curves conditioned on the (*n* − 1)th stimulus: fig. 4c shows that the PSE of the nth trial is pushed up to stronger stimuli after experiencing a strong stimulus on the (*n* − 1)th trial, and vice-versa. The same bias emerges in the RNN model as a consequence of Hebbian plasticity. High-intensity inputs shift the location of the attractor towards high intensity inputs, cf. fig. 3f. This shift translates to the effect shown in both measurements: a higher probability of categorising a given stimulus as weak (or strong), for a stimulus distribution biased towards higher (or lower) intensities, as shown in fig. 4f, and a shift in the PSE of the psychometric curves, given the different (*n* − 1)th stimuli (fig. 4g). In appendix D we discuss the quantitative disparity between fig. 4b versus f, which appears to stem from individual subject differences –a phenomenon we can also replicate with different random realizations. Here we emphasise that the results presented reveal that the model captures, in general, the relative importance of the preceding stimuli on the choices, as reported in fig. 4d and h.

### The one-back experimental paradigm – plasticity and relaxation at a glance

Our analysis highlighted the importance of the movement of the attractor in the RNN for recency and repulsive biases to emerge in perception. To challenge the generality of our model, we designed a novel **one-back task**. Participants undertake a sequence of trials, each of which presents a single tactile stimulus, similar to the reference memory task. However, triggered by the go cue, they must compare the current trial’s stimulus not to a fixed category boundary but to the stimulus of the previous trial. Comparing the current stimulus to the preceding stimulus is akin to the working memory task, however there are marked distinctions. First, the two stimuli are not bundled together in one trial but are distributed to sequential trials. Second, each stimulus has two functions: it is initially the comparison stimulus for judgment relative to the preceding trial but, with choice made, it now becomes the base stimulus for the next trial (fig. 5a). A crucial feature of the paradigm that tests for the dynamics of the attractor is that stimulus intensities are grouped into two clouds, a “low” intensity cloud with stimuli labelled from 1 − 4 and a “high” cloud of stimuli labelled from 6 − 9. After a stimulus is drawn from the low/high cloud, the next stimulus is drawn either from the same cloud or is the intermediate stimulus, 5. A transition from one to the other cloud may occur only with a trial at the intermediate stimulus. The transition probabilities *p*(*S*^(*n*)^|*S*^(*n*−*1*)^), shown in fig. 5a, can lead to long sequences of stimuli constrained to one of the clouds (mean within-cloud string length is 18 trials; median is 13). This paradigm tests the prediction that subjects’ psychophysical results will reflect the attractor’s continuous movement between the two clouds, as the trace of past stimuli transitions between “low” and “high.”

**Figure 5:**
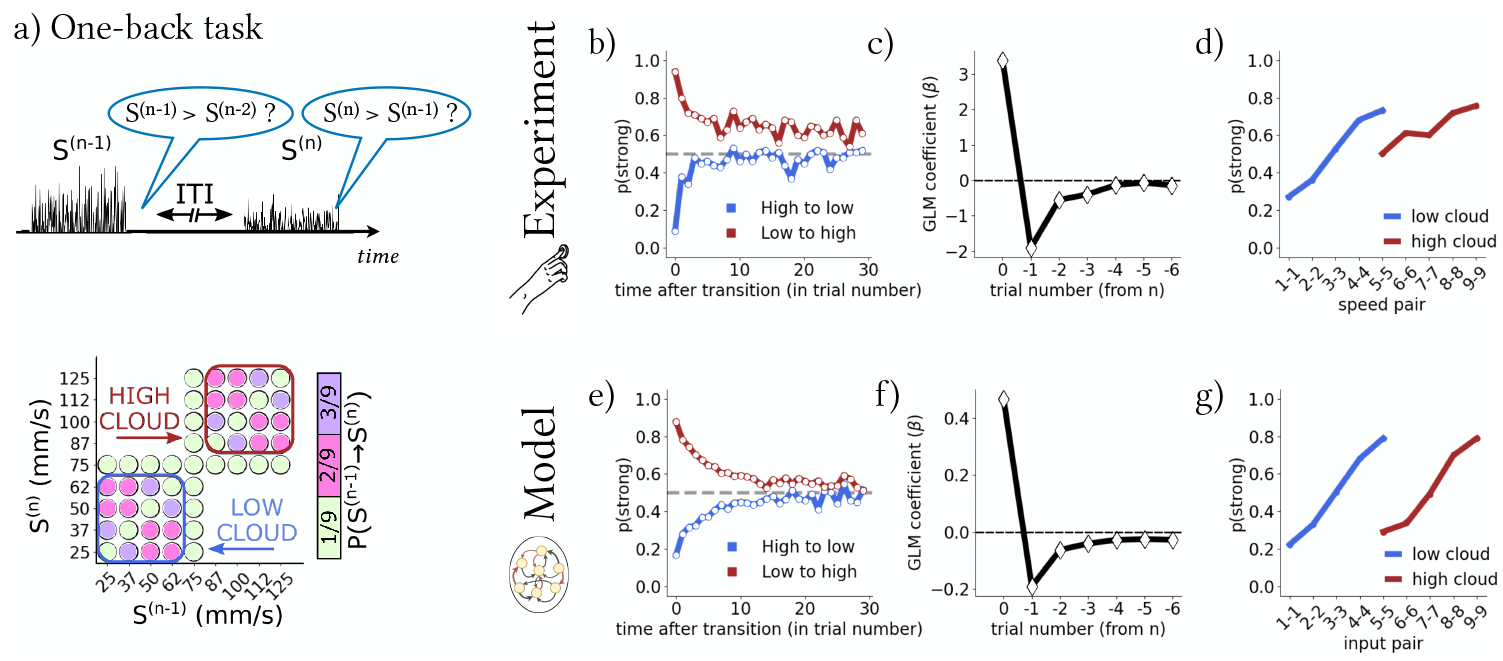
The one-back task highlights the plasticity of the attractor as it moves between stimulus clouds. **a)** We introduce the novel “one-back” task, where subjects are tasked to compare the strength of one stimulus per trial to the strength of the stimulus in the preceding trial (top). We design the transition probabilities governing the strength of the nth stimulus given the strength of the (*n* − 1)th stimulus for this task (bottom) such that long consecutive sequences of stimuli from the low-intensity or high-intensity “clouds” appear. **b)** Probability of human subjects categorising a stimulus as stronger than the preceding stimulus within each cloud as a function of the time elapsed since the transition between clouds. **c)** Coefficient values (*β*) of the generalised linear model (GLM) of the choice at trial n given the six preceding stimuli. **d)** Probability of categorising *S*^(*n*)^ > *S*^(*n*−1)^ when *S*^(*n*)^ = *S*^(*n*−1)^. If the pair 5, 5 is preceded and followed by stimuli from the low cloud, it is considered part of the low cloud; and vice versa for the high cloud. **e–g)** Results obtained from the RNN model.

The experiment comprised twenty-four human participants, each completing 750 trials. Figure 5b shows the probability that a given stimulus was judged as stronger than the previous as a function of time after a jump between clouds. By objective physical measures, within each cloud the (*n*)th stimulus is equally likely to be weaker or stronger than the (*n* − 1)th stimulus. But perceptual biases were uncovered in actual participants. After a switch from the high-to the low-intensity cloud, participants consistently underestimated the strength of the (*n*)th stimulus (against the (*n* − 1)th stimulus) over the first few trials (blue line), before stabilising around the correct value of 0.5. Conversely, after switching from low-to high-intensity stimuli, stimulus intensities were overestimated initially (red). The RNN model generates analogous behaviour ((fig. 5e). The model’s results are explained by the attractor migrating gradually, after the cloud switch, from its previous position around the average intensity of the opposite cloud. As the (*n* − 1)th stimulus representation is pulled towards the earlier cloud, a strong bias produces the observed misclassifications. As the stimulus string remains in the current cloud, the attractor slowly settles to a new globally stable position with small dynamical movements around the average intensity value of the cloud.

Averaging over time, the overall influence of previous stimuli on the choice can be estimated in terms of GLM coefficients as done in the previous two paradigms, showing similar results: the choice at trial n depends with a decaying weight on the previous stimuli as found in both experimental and model data in fig. 5c) and f).

The one-back paradigm highlights that the shifting of internal representations, which is responsible for contraction bias, does not occur towards the global average of stimulus intensities, which would be the centre of the stimulus range, intensity 5; instead, internal representations contract towards an intensity value that changes in time. The contraction bias in the one-back task is most evident considering choices of participants after experiencing the same input successively, *S*^(*n*)^ = *S*^(*n*−*1*)^. The probability of categorising *S*^(*n*)^ *> S*^(*n*−*1*)^ deviates from 0.5 in a discontinuous trend, shown in fig. 5d for human subjects. For instance, within a stream of low-intensity stimuli, the second presentation of stimulus of intensity 5 is consistently categorised as stronger than the previous stimulus of intensity 5, while the opposite is true within a stream of high-intensity stimuli. The internal representation of intensity 5 contracts towards a *lower* value when it is part of a low-intensity sequence, and vice versa, consistently biasing the categorisation of the second instance of intensity 5. The RNN model fully reproduces this behaviour (fig. 5g). Indeed, during the ITI the internal representation of the stimulus relaxes towards the running attractor, replicating the temporal dynamics of actual human subjects. Note that the asymmetry in the experimental results for stimuli from the high vs. the low cloud in fig. 5d, and the decay of the red curve to a value larger than 0.5 in fig. 5b, arise from a combination of the Weber-Fechner law and the stochasticity in the stimulation setup. In our model, where the Weber-Fechner effect is not included, the asymmetries in fig. 5e and g stem solely from this stochasticity in the stimulation, as detailed in appendix E. We emphasize, again, that this simulation was achieved with no external re-setting of the hyper-parameters *γ, λ, η, g* used previously for working memory and reference memory simulations.

## Discussion

Perceptual biases [3, 5, 7–12] offer a window into the computational processes underlying perceptual judgements, but the neural mechanisms from which they emerge remained poorly understood. Here, we proposed a simple recurrent neural network model with ongoing Hebbian plasticity that reproduces perceptual biases across four datasets spanning three different experimental paradigms, without any per-paradigm fine-tuning. Combining Hebbian plasticity with a set of biologically inspired input patterns yields a dynamic point attractor. Contraction arises from relaxation towards this plastic attractor, and the movement of the attractor, driven by Hebbian plasticity, leads to both repulsion and recency effects.

Our newly designed “one-back” task accentuates the movement of this plastic attractor, and the RNN model correctly predicted the novel bimodal contraction bias we observed in the performance of human subjects on the task.

Figure 6 provides a closer look at the time traces of the neurons in our model. We report the intensity values of the internal representation (gray) and of the attractor (green) in on sample simulations per paradigm. While the dynamics of neurons and weights follow the same dynamical equations (1) and (2) in all three settings, it is the task-specific input distribution that leads to different choice statistics. Wrong choices, highlighted by red rectangles, reproduce the task-specific perceptual biases.

**Figure 6:**
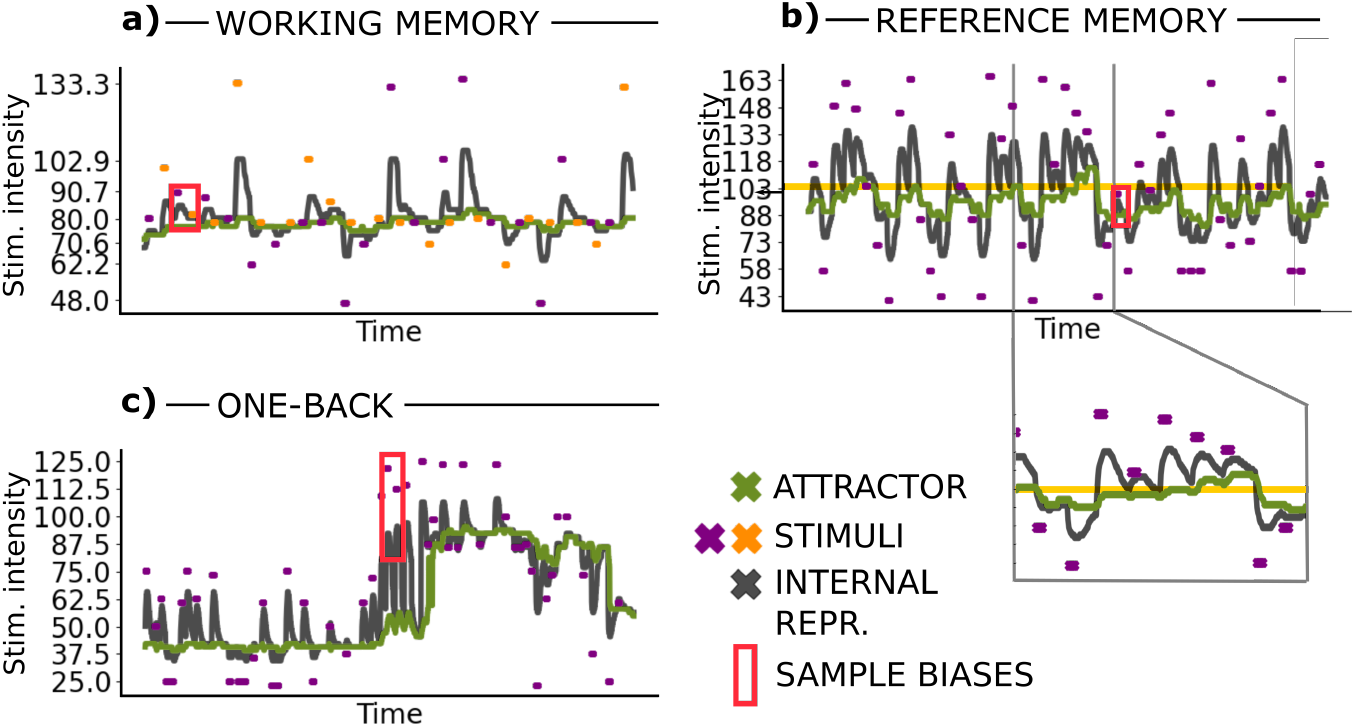
One neural network reproduces two different biases across three different paradigms: Evolution of the network in a sample working memory **a)**, reference memory **b)**, one-back **c)** task. The external stimulus is depicted in violet (and in orange when corresponding to *S*_*B*_ in **b)**, the internal representation (dark gray) and attractor position (green) at each time step is readout through the maximum of the overlap profile. The red rectangles indicate instances of incorrect choices while the orange horizontal line in **b)** represent the overall average.

There are two important timescales in our model. One governs the relaxation of the internal representation in the absence of stimulation, while another governs the movement of the plastic attractor. While the relaxation occurs on a shorter timescale than the movement of the attractor, the two timescales are comparable, and both evolve in time. The movement of the attractor’s location, which we refer to as the trace of past stimuli, depends on the states of the RNN 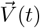, which in turn depend on the attractor itself and on the external stimuli. These timescales are thus only indirectly fixed by the hyper-parameters of the model; instead, the memory decay of a stimulus representation occurs over an *effective time scale* which emerges from the interaction of individual neurons and from the continuously evolving connectivity. We show in fig. F.4 that this time scale is indeed much longer than the timescale of individual neurons *γ*, meaning our model addresses the key challenge of modelling working memory, namely overcoming the discrepancy between the timescale of the dynamics of individual neurons and the timescale of memory [18]. This is particularly evident in the one-back paradigm sample reported in Figure 6c: a given stimulus influences the speed of movement and the effective value of both the internal representation and the attractor differently, depending on whether it occurs at the beginning or end of a period of continuous stimulation with stimuli from one of the two clouds. A promising direction for future research would be to explore how our findings relate to previous studies suggesting that the perception of probabilities extends beyond the running average [35, 36].

The idea that perceptual biases may arise from attractive dynamics has long been hypothesized. Initially, this concept was framed through a phenomenological model involving two interacting attractors with different timescales [10, 15, 16]. More recently, Boboeva *et al*. [20] incorporated this idea into a computational model that postulates two static continuous attractor neural networks with pre-determined connectivity and tuning curves to reproduce a single experiment. In our model, we let a single recurrent neural network learn its connectivity from stimuli via a local, biologically inspired learning rule, and obtain a model that generalizes across behavioural tasks and species. While in Boboeva *et al*. [20] contraction bias results from the difference in time-scale of the two networks and the presence of adaptation only in one of them, in our model contraction bias is an emergent effect of learning.

Our model also successfully reproduces a key result that was previously considered evidence for the concept that perceptual biases arise from the interaction between two neural networks: in a study by Akrami *et al*. [3], inactivation of the PPC subregion was shown to reduce contraction bias in an auditory working memory tasks, despite no loss of memory capabilities. We found that our RNN model replicates this behaviour following PPC inactivation by adjusting synaptic homeostasis, as shown in fig. C.1. Modifying the synpatic homeostatis effectively places the model in a regime in which the attractor tends to remain stuck at the intensity of the last perceived stimulus. This adjustment preserves memory capacity while reducing contraction bias, as the relaxation dynamics only minimally varies the effective coded intensity. Our findings suggest an alternative role for the PPC sub-network in the modulation of contraction bias.

An intriguing direction for future research is to confront our RNN model with neural recordings. In particular, recordings from high-density electrophysiology in brain regions with recurrent circuitry would lend themselves ideally for a comparison to our high-dimensional neural network model. To start with, it would be interesting to look for signatures of a dynamically moving attractor in neural recordings from animals performing perceptual tasks. This could be achieved by analysing how input intensities are represented by the neural population, followed by monitoring the relaxation of neural activity once the stimulus is removed. In the future, it will also be interesting to contrast neural representations and their geometry [37–39] found *in vivo* with those obtained from RNN models whose connectivity was shaped by Hebbian plasticity, as in our case, and RNNs trained with gradient descent.

Our model has some limitations. For simplicity, we have assumed a dense network with all-to-all recurrent connections, although it is unclear whether this accurately reflects the connectivity in VM1. Additionally, our model assumes that inputs predominantly originate from a single region, with a balanced input distribution (50% code positively, 50% code negatively external input intensity). However, recordings from VS1 suggest that the majority of units actually code intensity positively. Future work could investigate the effects of sparser connectivity and unbalanced input schemes on the network’s behaviour.

Our model shows how attractors, which have long been contemplated to code intensity in the brain [10, 19, 20], can be learnt continuously from the input distribution using a local learning rule. Our study also highlights the potential significance of transient RNN dynamics for encoding and processing information. The mathematical analysis of an RNN with ongoing Hebbian plasticity driven by intermittent, correlated inputs is a significant challenge. The recent dynamical mean-field theory for coupled neuronal-synaptic dynamics by Clark & Abbott [40] represents an excellent starting point. Finally, extending this model beyond sensory perception is another promising direction for further research.

## METHODS

### EXPERIMENTAL MODEL AND STUDY PARTICIPANT DETAILS

#### Experimental protocol for working memory task

Sixteen subjects participated in the WM task. All subjects performed one WM session, lasting approximately one and a half hour for a total of 1000 trials. All subjects were volunteers and were paid after the participation, on the basis of how well they performed. The study protocol conformed to international norms and was approved by the Ethics Committee of the International School for Advanced Studies (SISSA).

All experiments took place in the SENSEx Lab at SISSA. The subjects sat in front of a PC screen with their right arm resting on a pillow upon the desk, using their right index fingertip to trigger an infrared sensor in order to begin a trial; when doing so, their fingertip would be in contact with a plastic probe. The probe was attached to a shaker motor (type 4808; Bruel and Kjaer), producing vibrations in the horizontal direction, perpendicular to the fingertip. Subjects’ responses were reported by pressing one of two buttons with the right hand. The experiment was automatised via LabVIEW (National Instruments).

Each stimulus was a noisy vibration, obtained by stringing together randomly sampled velocities. Velocities were obtained by sampling a normal distribution with 0 mean and defined by the standard deviation *σ*, ranging from 3 to 20; the mean stimulus speed is directly proportional to *σ*, and the motor amplifier gain was set so that the average stimulus speed would be 10 times the value of *σ*, measured in mm/s. The sequence of velocities formed one seed. There were 50 different seeds for each vibration mean speed, making the signature of a given vibration hard to recognise and making the mean speed a more salient feature.

The subject’s finger had to remain in the sensor for the entire trial, comprising: pre-stimulus delay (500ms), base stimulus (334ms), inter-stimulus-interval (either 2, 4 or 8s), comparison stimulus (334ms), post-stimulus delay (500ms); after this sequence elapsed, a visual go cue signalled to the subject to report their choice. If the subject lost contact with the sensor before the go cue, the trial aborted, and a new one had to be initiated.

Nominal pair difficulty does not depend on the value of the stimuli, but on the relative difference between them: the closer they are, the more difficult the discrimination. We expressed this difficulty using an index, the Normalised Speed Difference (NSD), according to Weber’s Law:

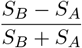

The stimuli pairs were arranged in two sets: a vertical set, in which the *S*_*A*_ *σ* was fixed at 8, and a horizontal set, in which the *S*_*B*_ stimulus *σ* was fixed at 8. For both sets, NSD values ranged from −0.25 to 0.25, including 0, and the non-fixed *σ* was calculated accordingly. Stimuli pairs from both sets were randomly presented during the WM sessions.

The GLM model employed a logistic link function and the parameters were estimated via Maximum Likelihood Estimation by bootstrapping trials on a subject by subject basis.

#### Experimental protocol for one-back task

Twenty-four human subjects (ages 19-38), recruited from the online research participation system (SONA), participated in the One Back Experiment. The study adhered to frame of rules specified by international norms for human behaviour experiments and was approved by the Ethics Committee of the International School for Advanced Studies (SISSA).

The experiment was conducted in Human labs at SENSEx Lab in SISSA. The subjects were seated in a dimly lit room. The experiment was executed in LabView software. Subjects viewed a monitor screen for cues and feedback related to the task. They wore headphones that presented white noise and eliminated any external sounds from the environment or motor. The vibrotactile stimuli were delivered to the tip of their right-hand index finger through a shaker motor (type 4808; Bruel and Kiaer). The responses were recorded using keyboard button press using left hand. The subjects saw a blue dot as fixation on the screen. They received feedback (green/red dot for correct /incorrect) on each trial through the screen.

The subjects performed a one-back memory task where the goal was to compare stimulus *S*^(*n*)^ with *S*^(*n*−*1*)^. The stimuli were generated by sampling a sequence of velocity values. The probability of velocity values was drawn from a normal distribution (*μ* = 0, SD = *σ* in mm/s). The gain of the motor was amplified 10 times. The velocity sequence was taken randomly from 50 seed values for a given trial. These were transmitted as voltage values to the shaker motor producing motion in horizontal direction using a plastic knob delivered perpendicular to the finger. The stimuli had a fixed duration of 350 ms. The judgement is based on the mean speed of the stimulus, which is subjectively reported by subjects to be “strength”, “amplitude”, or “intensity”. The mean speed of the vibration was proportional to the SD values used. Nine SD values [2.5, 3.75, 5, 6.25, 7.5, 8.75, 10, 11.25 and 12.5] were used.

Each trial began with stimulus delivery followed by response time window (timed on the screen by a graphical motion of blue colour filling a vertical bar) followed by Inter-Trial Interval (ITI). The subjects had two seconds available to report their decision by pressing keys [A or D]. After every response, the feedback was displayed on the screen. Failure to respond within the given response time resulted in a timeout. Three ITI values [0, 2 and 6 sec] were used, resulting in total ITI of 2, 4 or 8 seconds between two consecutive stimuli. Reward was delivered for correct choices (if the same stimulus intensity is presented on two consecutive trials, the correctness of the comparison is rewarded randomly.)

The experiment lasted one hour and was divided into 3 blocks (250 trials each). Each subject performed a total of 750 trials. The average performance of subjects in One Back task is 76.63 percent. All data analyses were performed using custom scripts in MATLAB. The GLM model was calculated as introduced in the previous section.

## METHOD DETAILS

### RNN model

#### Motivation

The four psychophysical experiments that we reproduce with our model – a reference memory, two working memory, and a one-back memory tasks– are comprised of judgements made on vibratory stimuli delivered to the fingertip of human participants, to the whiskers of the rats or by auditory stimuli delivered to rats. As far as rats and tactile stimuli are considered, the model may be considered a representation of the frontal cortical region known as vibrissal motor cortex (vM1), the main target of somatosensory cortex in rats [41]. vM1 is involved in memory, decision making, and motor planning and is a recurrent network that receives afferent inputs from the primary vibrissal somatosensory cortex (VS1) [41]. In humans, the corresponding cortical regions that fulfil these tasks are currently unknown, though new evidence is emerging [42].

#### Sources of Randomness

Different realizations of the model arise from several sources of randomness: a) The realization of the input pattern scheme; b) The realization of the stimulation protocol; c) The stochasticity in the stimulation setup for each delivered input; d) The random initialization of the weights. The random protocols for a, b, and c are described in appendix A), the one for d in the following *Simulation* subsection.

### Input patterns

Our model for the afferent inputs to the RNN units is inspired by physiological recordings from Arabzadeh *et al*. [30] and Fassihi *et al*. [31] of the primary vibrissal somatosensory cortex (VS1). They found that the majority of vS1 neurons in behaving rats show monotonically increasing firing rates as vibration intensity increases, with heterogeneity across neurons in regard to sensitivity (minimum vibration to cause excitation) and slope. We set the ratio of positive versus negative slopes to 50% to account for the possible presence of interneurons. Half of the neurons receive inputs whose magnitude increases with stimulus strength (positive slope), while the other half receives inputs that decreases with stimulus strength. We discretize the intensity line, from a minimum to a maximal value, into P bins labelled *μ* = 1‥*P*. We thus obtain an input pattern matrix 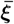 of size *N* × *P*, where 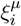 is the input unit i receives when the intensity value corresponding to the *μ* label is delivered to the network. We provide additional details about the generation of *input patterns* used in our model in appendix A. In addition, we also model the stochasticity of the experimental stimulation setup, a minor detail also explained in appendix A, for which, essentially, each stimulus strength is actually defined by a standard deviation. The specific values for the standard deviations used to sample vibrational stimulus strengths and their respective extraction probabilities used in both the tactile experimental sessions and the model are as follows:

#### Working Memory Task

Standard deviation values (*σ*) were 4.8, 6.22222, 7.05888, 8, 9.06667, 10.28577, and 13.33333. The *S*_*A*_ and *S*_*B*_ couples were organised as depicted in fig. 2a)e), with each pair extracted with equal probability.

#### Reference Memory Task

Standard deviation values (*σ*) were 4.3, 5.8, 7.3, 8.8, 10.3, 11.8, 13.3, and 14.8. Each value was extracted with equal probability, except in fig. 4 b)f), where biased extraction probabilities were indicated in the plot.

#### One-Back Task

Standard deviation values (*σ*) were 2.5, 3.75, 5, 6.25, 7.5, 8.75, 10, 11.25, and 12.5. Extraction probabilities were as reported in fig. 5).

### Simulations

All simulations were conducted in Python, and we provide code to run them online. We employed the following hyperparameters for eqs. (1) and (2): *γ* = 0.1, *λ* = 0.014, *η* = 2, and g = 1.2, *ϰ* = N, facilitating a homeostatic regime where learning and dynamic evolution proceed continuously without stagnation. For simplicity, we set the limiting average activity a, towards which the network’s average activity is driven by the threshold, to match the average input activity. This value depends on the random realization of the input patterns in each simulation and is given by 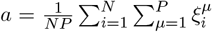, typically yielding approximately 0.2. Variations of these parameters are also feasible. We simulated an RNN with *N* = 200 units and parametrise the spectrum of weak to strong vibrational stimuli with *P* = 100 input patterns. We initialised the RNN weight matrix with random weights ranging from −1 to +1 at the beginning of each “experimental session”, i.e. at the beginning of each run. Averages where performed on 100 random realisations each composed by 500 trials for the working memory paradigm, 200 random realisations each composed by 300 trials for the reference memory paradigm and 150 random realisations each composed of 750 trials for the one-back task. The temporal parameters of the experimental protocol are configured as follows.

#### Working Memory Task

5 time steps before each stimulus, 5 time steps for stimulus presentation, followed by randomly selected intervals of 5, 10, or 20 time steps before the second 5-time step stimulus, followed by a randomise interval of 10-15 time steps before the following trial.

#### Reference Memory Task

10 time steps before the stimulus, 10 time steps for stimulus presentation, and a randomised interval of between 10 to 20 time steps after the stimulus.

#### One-Back Task

10-time steps for stimulus presentation, followed by randomly selected intervals of 20, 40, or 80-time steps from one stimulus to the next. In figs. C.1 and E.1 we used a different stimulation protocol as reported in the capitation.

Note that the exact relationship between these time steps and real milliseconds is beyond the scope of this article. However, we demonstrate in appendix F that this specific choice is not a fundamental feature to obtain the results shown in the article.

#### Reading out the network

We read out the intensity which is internally represented by the RNN model at each time as the intensity value corresponding to the stimulus that has the largest cosine similarity, or overlap, with the RNN state 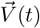. We perform the choices based on these representations as follows. In the working memory and in the one back paradigms, we compare the internal representation to the delivered stimulus at the stimulus onset (only for *S*_*B*_ in working memory). If the intensity of the delivered stimulus is larger than the internal representation of the network, the delivered stimulus is considered stronger than the previous one, and vice versa. In the reference memory paradigm, we consider the relaxation dynamics of the internal representation at the stimulus offset. If the internal representation relaxes towards higher intensities by the end of the trial, the stimulus is considered weak, and vice versa. When the choice is ambiguous, i.e. if the (second) stimulus coincides with the internal representation for working memory and one-back, or if the internal representation does not relax in reference memory, we take a random choice. Furthermore, we introduce “lapses” in all three paradigms, which reflect the animal’s tendency to make errors due to a lack of concentration. Specifically, we take at random the choice in 10% of the trials, which are picked at random.

#### Fitting psychometric curves to data

We fitted the psychometric curves in the experimental and the simulated data of figs. 2, 4 and C.1 using a the same modified sigmoidal function as Hachen *et al*. [10], which has the form

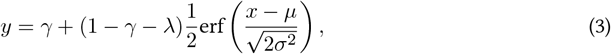

where *γ, λ, μ*, and *σ* are all fitted to data (see Hachen *et al*. [10] for a detailed discussion of the motivation and interpretation of the fitting parameters). In figs. 2 and C.1, *y* is the probability that the second stimulus is judged stronger than the first, *S*_*B*_ *> S*_*A*_, and *x* is a categorical variable indicating the difference between stimulus intensity. In fig. 4, y is the probability that a given stimulus is judged “strong”, and x is a categorical variable indicating the stimulus label with the weakest stimulus corresponding to 0.

## Data Availability

The code used to simulate the RNN is available at osf.io/msbuh.

## Acknowledgements

We thank Iacopo Hachen for valuable discussions and for providing the data used in fig. 4. We thank Alessandro Treves for valuable discussions on Blumenfeld *et al*. [32]. F.S. is supported by the QBio Junior Research Chair program of the QBio initiative of ENS-PSL. SG gratefully acknowledges funding from the European Research Council (ERC) for the project “beyond2” under the European Union’s Horizon 2020 research and innovation programme, Grant agreement ID 101166056, and co-funding from Next Generation EU, in the context of the National Recovery and Resilience Plan, Investment PE1 – Project FAIR “Future Artificial Intelligence Research”. This resource was co-financed by the Next Generation EU [DM 1555 del 11.10.22].

## Author contributions

F.S, M.D. and S.G. designed the study. F.S. proposed the theoretical model, developed the simulations and performed the experimental-theoretical comparisons. D.G. performed the experiments on the Working Memory paradigm and Y.C. performed the experiments on the One-Back paradigm. F.S. and S.G. wrote the first draft of the paper and F.S., S.G. and M.D. carried out subsequent revisions. All authors approve the final manuscript.

## Appendix

### A Input patterns, stimulation protocol and stimulation setup stochasticity

#### Input patterns

Here, we provide additional details about the generation of input patterns utilised in our model. The inputs are represented by a matrix 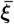 of size *N × P*, where *N* is the size of the network and *P* are the number of bins discretising the intensity values from a minimal to a maximal one. Each row 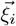 denotes the input profile received by neuron *i* for every intensity value. Each column 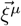 represents the pattern of activity delivered to the network when a specific input intensity, labelled as *μ*, is present, with 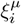 denoting the intensity input received by neuron *i* for input *μ*. To discretise intensity values, we divide a range from a minimum *s*^*MIN*^ to a maximum *s*^*MAX*^ intensity into P bins, each coding for the intensity ((*S*^*MAX*^ − *S*^*MIN*^)/*P*). To generate the input patterns, we employ the following procedure: we begin with a vector of length N containing random numbers ranging from −1 to +1; we create a second vector by replacing each negative entry in the first vector with a random number between 0 and 1, and each positive entry with a random number between −1 and 0; we linearly interpolate *P* − 2 patterns from the first to the last vector, ensuring continuity between intensity levels; finally, we set any negative values to zero to maintain non-negativity in the generated patterns. Through this process, we obtain a set of input patterns ranging from 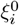 to 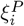. The pattern statistics can thus be summarised as follows: Approximately 50% of units receive positively coded inputs, with the remaining units receiving no input for each intensity (fig. A.1a, displaying 50 sample profiles 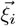 with different colors). The average input pattern activity exhibits a slight decay towards *ξ*^*P/*2^ before increasing again (fig. A.1b). The fraction of units receiving a non-zero input remains roughly constant across patterns (fig. A.1c).The correlation structure between patterns exhibits a monotonic shape, which explains why the dynamics observed in the network after progressive Hebbian learning can be likened to a bell shape when considering the overlap with the inputs. In fig. A.1d, we show, in a sample with *P* = 50, the overlap 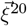 (light gray), 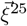 (black), and 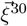 (dark gray) with all other patterns.

#### Stimulation protocol

The stimulation protocol in the four different tasks dictates the stimuli delivered and the probabilities of delivering each of them at every step, as in the experiments. In the model, at every step in time, the input *ξ*(*t*) appearing in eq. (1) is defined as follows:

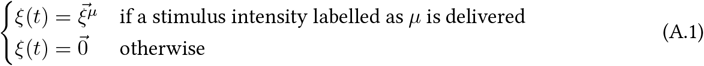

The precise time-steps defining the pre-stimulus interval, inter-stimulus interval (ISI), and inter-trial interval (ITI) for each paradigm are provided in the *Methods* section.

#### Stimulation setup stochasticity

For the tactile experiments, we also account for the additional variability introduced by the setup. While this is a minor detail, we include it to ensure a proper comparison between theory and experiment. As shown in the main text, the effect of this stochasticity is only mildly noticeable in the one-back paradigm, due to the very close-by intensities we selected.

In the experiments, as described in previous papers, see for example Hachen *et al*. [10], tactile inputs are delivered through a motor as a speed of vibration in a stochastic manner. A stimulus intensity is measured in terms of “average speed” 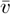. Each “average speed” results from a certain standard deviation *σ*. In fact, every 0.1 ms (10 kHz), a velocity drawn from a Gaussian distribution | 𝒢 (0, *σ*) |, where | |. is the absolute value, is delivered to the subject. The Gaussian distribution is illustrated for the set of stimuli delivered in the reference memory paradigm studied in Hachen *et al*. [10] in fig. A.1e. Negative values are converted to positive ones, resulting in a distribution with an effective mean 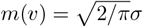 (vertical line in fig. A.1 f). Through a motor-gain parameter y, the effective average velocity is then converted to 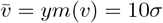. Basically, in the experiments, any time a certain intensity is delivered to the network, this may be a slightly different average speed (i.e. intensity) from one time to the other. We reproduce, in fig. A.1g, a sample distribution of the actual effective average velocities delivered across the different trials in a putative session of the published dataset in Hachen *et al*. [10].

**Figure A.1:**
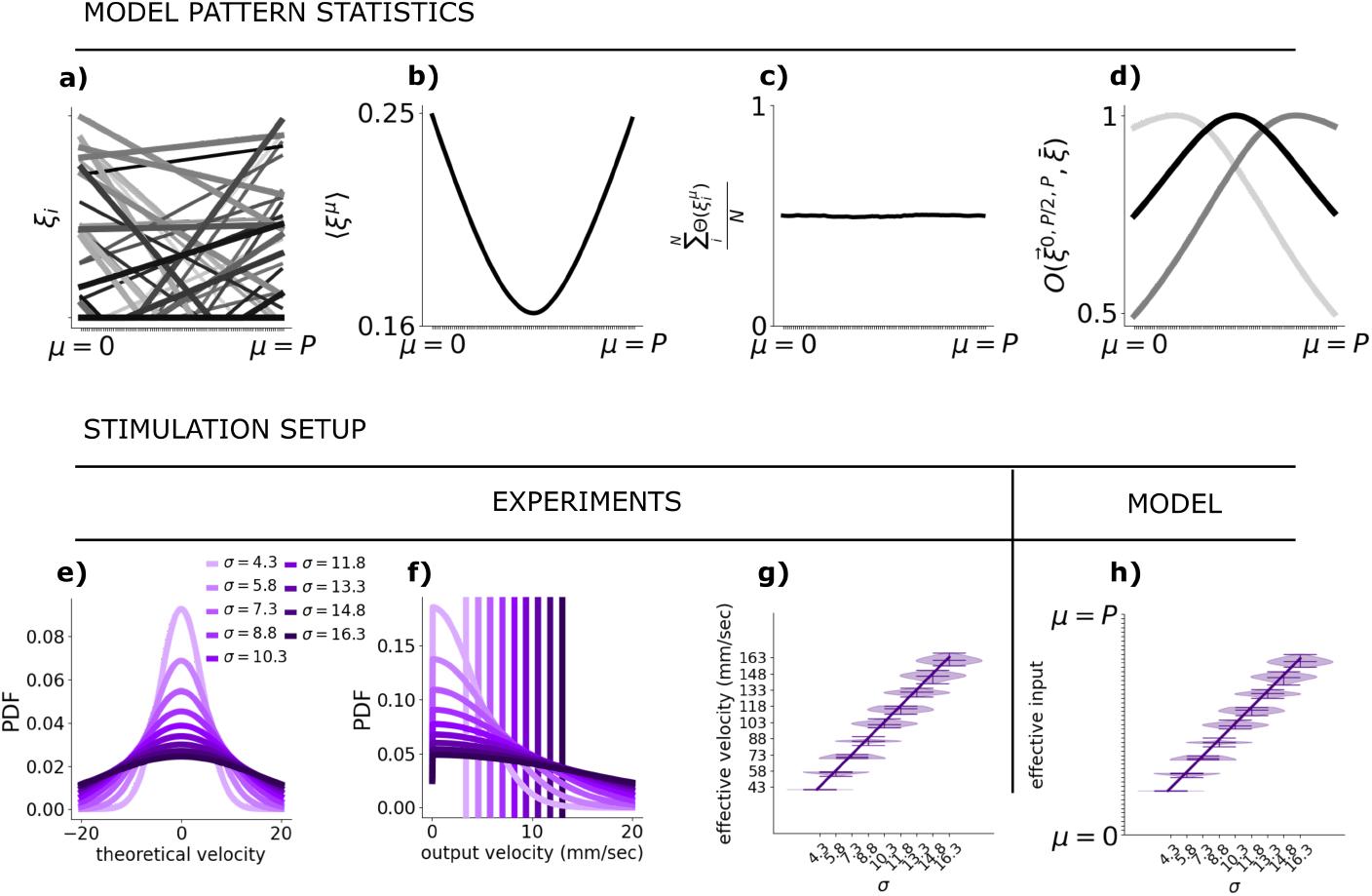
Input Pattern Statistics and Stimulation Setup Stochasticity: **a)** Fifty sample input profiles 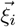 (different colours) depicted for different intensities. **b)** Average activity across input profiles averaged over 100 realisations. **c)** Fraction of active units in the input profiles averaged over 100 realisations. **d)** Normalised overlap (cosine similarity) between 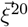 (light grey), 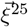 (black), 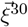 (dark grey), and all other patterns. **e)** Gaussian distribution with mean zero and standard deviation as labelled. **f)** Same as **e)**, but showing absolute values; the vertical line indicates the effective average activity. **g)** Violin plot representing the actual average obtained by sampling 5000 random numbers from the distributions in **f). h)** Identical to **g)**, but with the input intensities discretised into P bins, illustrating the transition between experiments and model.

In the model, we replicate the same bias in the distribution of the stimuli as follows: whenever an input defined by a standard deviation *σ* is delivered, we compute the average of 5000 random numbers drawn from the Gaussian distribution |𝒢 (0, *σ*) |. This average give the effective input intensity which is delivered. We pick the corresponding discretised input patter 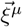 accordinglyThe quantity of 5000 was chosen to approximate the average intensity perceived over approximately 500ms.

In the figures we refer to the input intensity delivered (corresponding to the actual average velocity), and not to the standard deviations, in alignment with previous papers.

### B. Synaptic Weight Changes and Biological Plausibility

In the main text, eq. (2) consists of two terms: a homeostatic term and a standard Hebbian learning term. The homeostatic term (the first one) is crucial for preventing the divergence of synaptic weights during the learning process, thus ensuring the stability of the network. As a result, the weights can change from positive to negative over time. Although synaptic weights in the model are intended to represent the effective connectivity between neurons—including potential contributions from inter-neurons that add complexity—we believe it is still important to address the phenomenon of weight switching, which is in apparent contradiction with Dale’s principle. In this section, we provide a more detailed discussion of this behaviour and present preliminary analyses to clarify the role of these weight changes within our model.

**Figure B.1:**
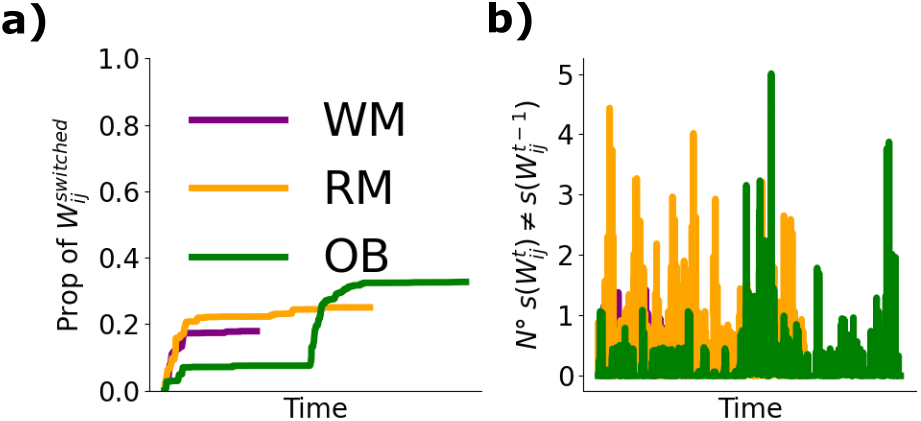
Excitatory/Inhibitory switch in the effective connectivityxs. **a)** Proportion of weights that have made at least one switch at the different stages of the trial and **b)** Absolute number of weights which have changed sign from one step to the other for the three examples shown in Fig. 6 of the main text. Specifically one per paradigm, labelled with different colour. Purple: Working memory (WM); Orange: Reference memory (RM); Green: One-back (OB). Please note that the time is non matching due to the different length of the trials. In **b** one should consider the absolute number of changes in comparison with the total amount of weights which is *N* ^2^, with *N* = 200

Figure B.1 presents an analysis of synaptic weight changes across the three example tasks shown in Fig. 6 of the main text. Our preliminary analysis reveals that approximately 20% of the total synaptic weights (as shown in fig. B.1 a) undergo at least one switch after a full trial of any paradigm. Interestingly, the maximum percentage of synaptic connections that change from one step to the next is 0.0125% (100 × 5/200^2^), as illustrated in fig. B.1 b. These weight switches are predominantly observed in synapses with low absolute weight values (not shown here), which are more likely to surpass the zero boundary during the homeostatic regulation process.

Recent studies have provided evidence for the presence of multi-transmitter neurons in the mammalian brain. For example, the work by Wallace & Sabatini [29] points to the existence of neurons that can switch between excitatory and inhibitory effects based on the relative presence of GABAergic and glutamatergic post synaptic receptors and transmitter received. This phenomenon, initially observed in the 1990s [43, 44], and also more recently explored by Morales & Margolis [45], suggests that some neurons may exhibit both excitatory and inhibitory activity under different circumstances. Nevertheless, introducing Dale’s law, which posits that each neuron typically releases only one type of neurotransmitter, in our model, is an interesting direction for future work, offering deeper insights into the dynamic behaviour of synaptic weights.

**Figure C.1:**
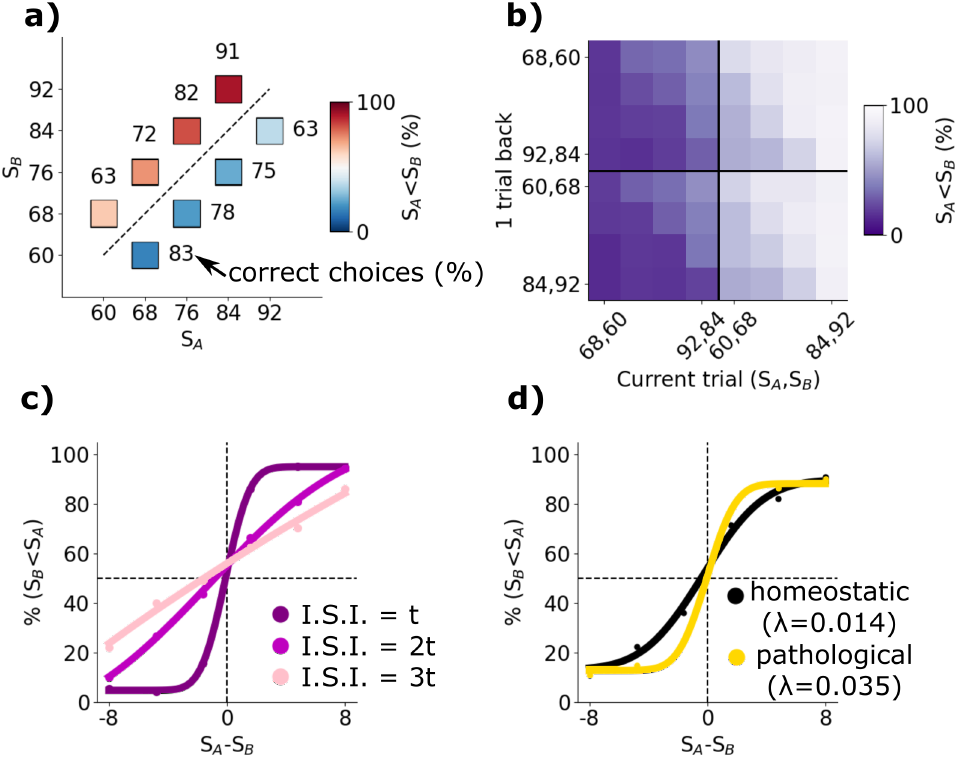
The RNN model replicates contraction bias and history-dependence demonstrated by Akrami *et al*. [3], see appendix C. **a)** Probability of correct choices for 8 stimulus pairs (*S*_*A*_, *S*_*B*_). Variations in the probability of correct choice among different pairs illustrate the presence of contraction bias, aligning with results depicted in Fig. 1b of Akrami *et al*. [3]. **b)** Probability of correct choice influenced by the stimulus pair presented in the previous trial, showcasing a trend where lower intensity pairs result in a higher probability of *S*_*B*_ being chosen stronger than *S*_*A*_. This mirrors the findings in Fig. 2a-c of Akrami *et al*. [3]. **c)** Performance across a subset of 6 stimulus pairs, each with a constant *S*_*B*_ = 76 and *S*_*A*_ ranging from 66 to 84. Results demonstrate the impact of ISI intervals on performance, with shorter intervals leading to better performance, consistent with Fig. 1e of Akrami *et al*. [3]. **d)** Performance variations with changes in the synaptic homeostasis parameter *λ*. Increasing *λ* induces higher performance by accelerating synaptic forgetting of previous stimuli and facilitating faster attractor updates. This pathological regime replicates effects observed during the optogenetic inactivation of PPC, akin to Fig. 3c of Akrami *et al*. [3]. Model parameters: Each stimulus is preceded by 5 time steps, followed by a presentation lasting 10 time steps for a)b) and 5 time steps for c)d). Subsequent to the initial stimulus, random intervals of 15, 30, or 45 time steps occur before the second 10 steps stimulus, with a random interval between 10-15 time steps between subsequent trials. Network dimensions: **a)** and **b)** N = 204, P = 102; **c)** and **d)** N = 196, P = 98. Hyperparameters *γ, λ, η*, and g remain consistent with the main model, except for D) (yellow), where *λ* = 0.035. Averages were performed over 50 random realisations each composed by 500 trials. In c) and d), points are experimental data and the solid lines are sigmoidal fits, see Methods.

### C Working memory paradigm of Akrami *et al*. [3]

We utilise the same model as described in the main text, including the same parameters, but with stimuli and inter-stimulus interval distributions similar to those presented in Akrami et al. (2018). Our simulations reproduce the observed contraction bias effects (fig. C.1a), the repulsive effect of previous stimulus pairs (fig. C.1b), and the dependence of contraction bias on inter-stimulus intervals (fig. C.1c) (to be compared with Fig. 1b, Fig. 2a-c, Fig. 1d of Akrami *et al*. [3] respectively). Furthermore, Akrami *et al*. [3] showed that inactivating the posterior parietal cortex (PPC) leads to a reduction in contraction bias while maintaining memory capability. In our model, which normally operates in a homeostatic regime where learning and transient neural evolution occur continuously, we observe a similar effect if we modify the synaptic homeostatic parameter *λ* in eq. (2). This adjustment leads to a regime where synapses quickly forget previous stimuli. As a result, the internal representation in the network tends to become stuck to the last perceived stimulus, which effectively becomes the new fixed point on average. The attractor is thus very closed to the last perceived stimulus and the drift of the memory trace during the ISI is minimal, consequently diminishing contraction bias while preserving memory until the next stimulus, see fig. C.1d (to be compared with Fig. 3c of Akrami *et al*. [3]).

**Figure D.1:**
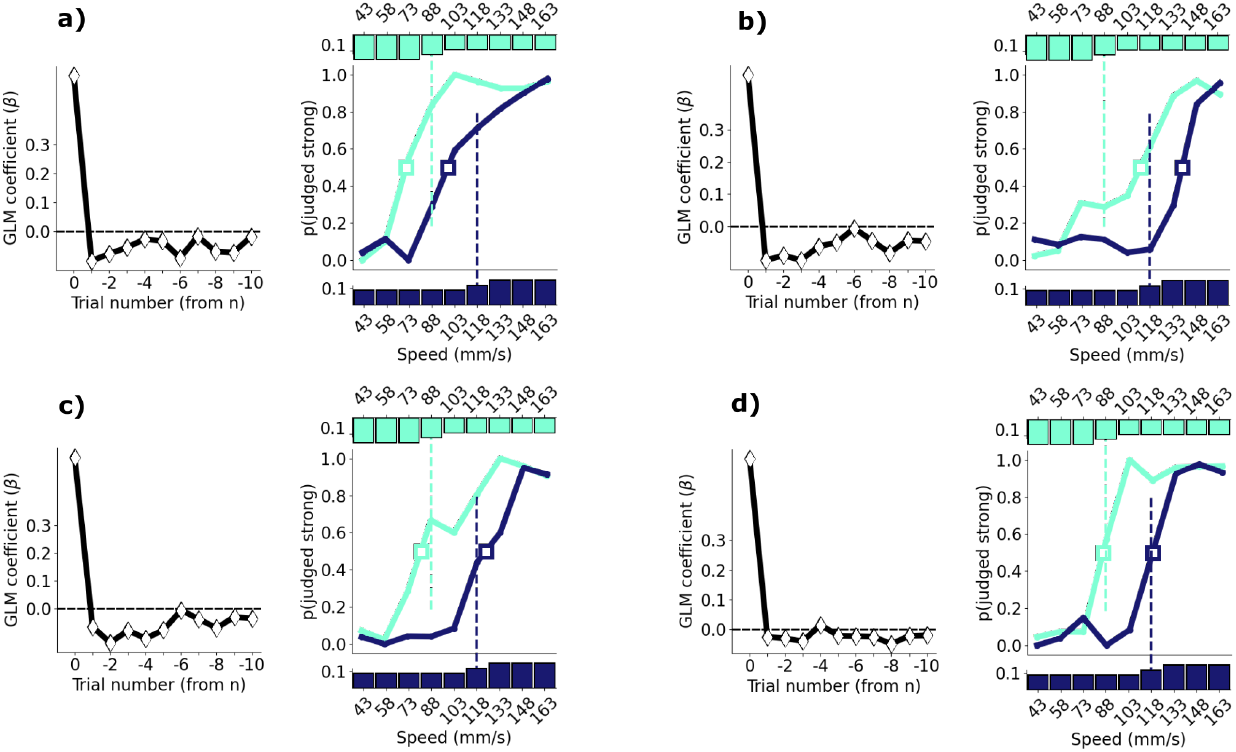
Individual differences between random realisations of Experiment 2. **a)–d)** Left, GLM as in fig. 4h. Right, psychometric curves as in fig. 4f (note that here the solid line is just joining the experimental points and is not a fit). Differently from fig. 4e–h we plot the results from 4 individual random realisations and not the average over 200 realisations.

### D Asymmetric shift of psychometric curves in Hachen *et al*. [10] and individual differences

Figure 4b in the main text reproduces a plot of Hachen *et al*. [10]. The reported shift in the psychometric curves is asymmetric: the psychometric curves obtained when more high stimuli were delivered during the session (dark blue) has a point of subjective equality (square) corresponding to the new average; instead, the point of subjective equality of the psychometric curves obtained from these sessions where more low stimuli were delivered (light blue), do not coincide to the new average and is actually even slightly shifted to the right as compared to the PSE for uniform distributions. On the contrary, the result of the model is a symmetric shift, as reported in fig. 4f. We asked further details to the authors of Hachen *et al*. [10]. First they reinforced that the major result reported by their figure is the effective separation between the psychometric curves, which we reproduce, more than the quantitative distance of this shift. Second, they reported that the seemingly absence of movement of the light blue psychometric curve, with respect to an hypothetical uniform distribution of stimuli, is attributed to the tendency of the tested subjects of underestimation, i.e. of psychometric curves with PSE shifted towards the right. Thus, it remain open the question on whether a larger subject pool would actually lead to a symmetric shift in the psychometric curves as that we predict through the model. Note that in the results of the model we use a large sample size, thus obtaining smooth results, but if we would focus only on a subset of seeds which give rise, by chance, to right-shifted psychometric curves, we would also reproduce an asymmetric shift. We can see individual differences in our model as different realisations of the model dynamics, which will differ from each other due to different random settings (see *Methods*). Randomness arise principally from the initial weights (akin to different subjects) and different random realisations of the input patterns and stimuli. In fig. D.1, we report four examples of the psychometric curves and GLM estimates for four different realisations, composing, together with other 196 realisations, the average values presented in the plots in fig. 4e–h. Individual peculiarities and stimulus history may lead to a different effects in the psychometric curves: for example in the realisation fig. D.1 a, there is a bias towards a strong categorisation, i.e. the attractor had a tendency to get stuck at high intensity, as opposed to the sample in fig. D.1 b.

**Figure E.1:**
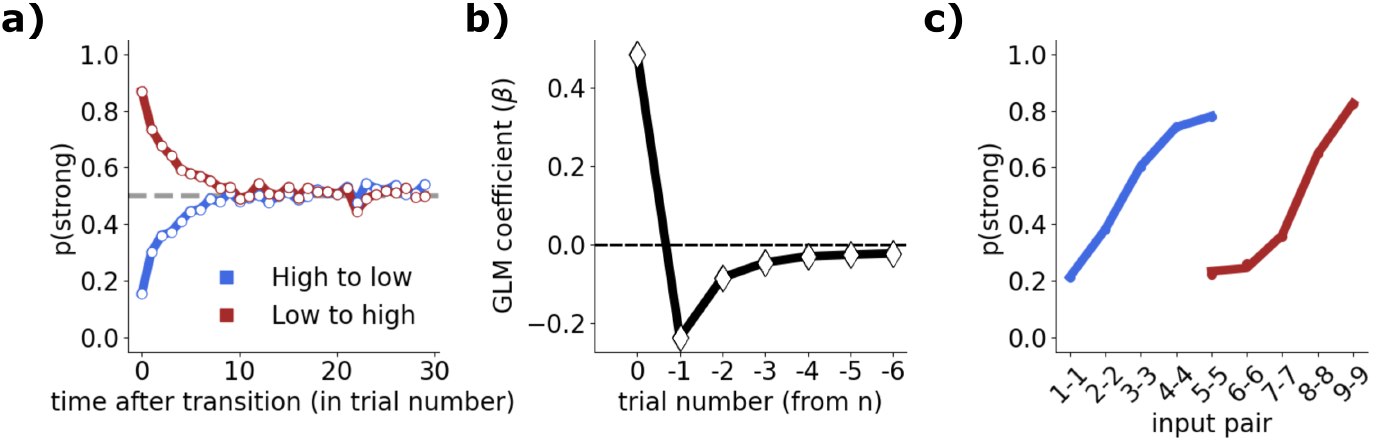
One-back Model without Input Stochasticity: The model and figures remain consistent with those reported in fig. 5e-g). The difference lies in the effective inputs, where the stochastic component of the stimulation setup is removed.

Finally, we hypothesise that if experimentally, even with a larger pool, the shift would keep being asymmetric another component of the model which could contribute in reproducing the behaviour is the distribution of the input pattern profiles. Indeed, in all results we presented so far, a ratio of 50% is considered between positively and negatively coding stimuli. Changing this percentage may have some effect of this type on the learned patterns.

### E The effect of the stochasticity in the stimulation setup

We note that the asymmetry in the results for stimuli from the high vs. the low cloud in fig. 5d, as well as the decay of the red curve to a value larger than 0.5 in fig. 5a, arise from two independent factors. Firstly, the Weber-Fechner law [46] which suggests that the subjective sensation of stimulus intensity is proportional to its logarithm. This phenomenon facilitates the comparison of two equally distanced quantities when they are small compared to when they are large. Secondly, the stochasticity inherent in the stimulation setup which generates a broader distribution of inputs when stimuli are high, cf. fig. A.1. In the model, but not in the experiments, we have the capability to eliminate the stochasticity inherent in the inputs, exposing the network solely to the average stimuli, thereby rectifying the asymmetries in the results, as shown in fig. E.1. Note that this effect is observed only in the one-back task because, in this design, the difference between consecutive intensities is much smaller than in the other experiments (see *Methods*).

### F Further details on time scales in the model

#### F.1 The ratio between inter-stimulus interval and stimulus duration

In the results presented in the main text, the ratio between stimulus duration and inter-stimulus interval (ISI) in our simulations was not consistent with the experiments. Here we show that our model also reproduces the contractive bias in the working memory task even when the ratio between stimulus duration and ISI matches the experiment. The top row of fig. F.1 reproduces the results from our simulation of the working memory task for convenience, as reported in the main paper. The bottom row of fig. F.1 reports the results of a simulation where we set the time step equal 333.5ms, resulting in 1 time step before each stimulus, 1 time step for stimulus presentation, followed by randomly selected intervals of 6, 12, or 24 time steps before the second 1-time step stimulus. All these durations precisely match the ratio of stimulus duration and ISI used in the experiments on real human subjects. Since the stimulus is applied for a much shorter time, we increased the stimulus intensity to *η* = 5 compared to the simulations reported in the main text.

**Figure F.1:**
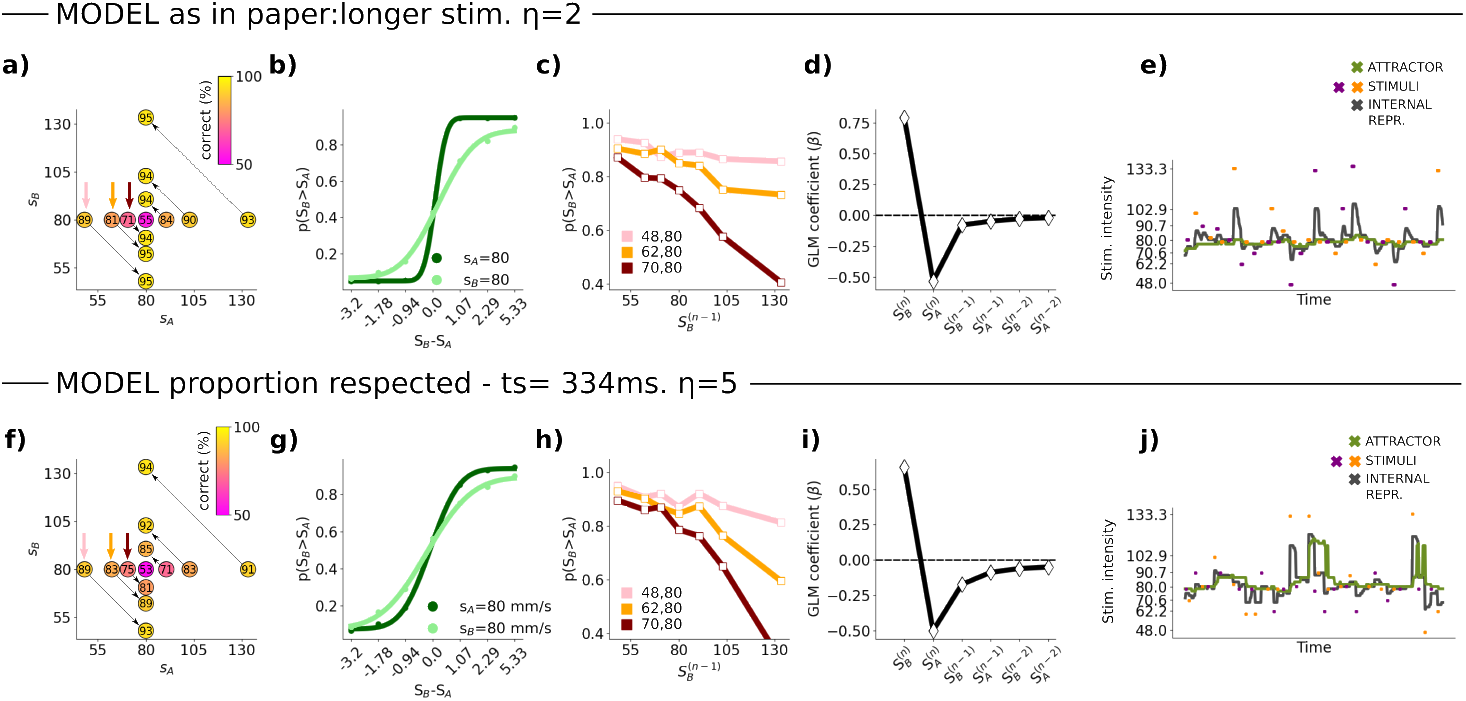
Our model reproduces contraction bias and history-dependence of choices in the working memory experiment when the ratio between stimulus duration and ISI in simulations matches the ratio in the experiment. **a–e)** reproduce the panels of Figs. 2 e–h and Fig. 6 a) of our manuscript and are obtained from simulations. **f–j)** show the same plots obtained in a simulation where the ratio between inter-stimulus interval and stimulus duration matches the experiment. We also increased the stimulus intensity to *η* = 5. Averages are taken over 100 simulations, each consisting of 500 trials.

In the main text, we report results from simulations with ratios that do not closely match experiments for computational efficiency: applying the stimulus for longer, with a smaller *η*, allowed us to perform trials in less total timesteps while having a stimulus duration that was long enough to track the dynamics of its effect on the attractor. We made this choice based on experimental evidence from Toso *et al*. [47], who showed that the percept of a stimulus is independent of its duration for a stimulation length above 300 to 500ms, see fig. F.2. We therefore assumed that we could extend the stimulus duration in our simulations without changing how the modelled subject would have experienced its intensity. As we show in the next subsection, our model reproduces other behavioural effects related to timescales with the hyperparameter settings used in the main text.

**Figure F.2:**
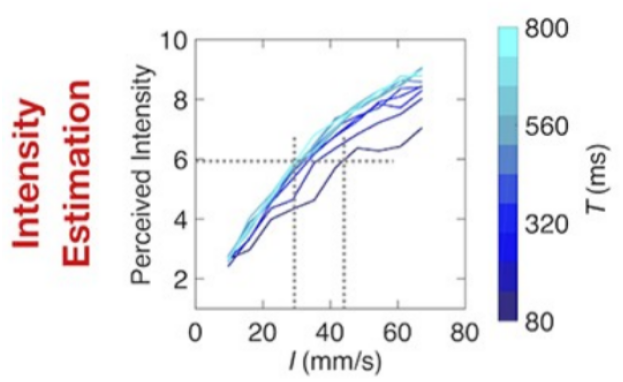
The perceived intensity of a stimulus depends on its duration *T* only for short *T*. Estimated intensity (ordinate) of vibrational stimuli as a function of their actual intensity (abscissa) and the stimulus duration (in different colours). The dense packing of curves for T greater than about 300 ms means that duration does not affect perceived intensity. Figure reproduced from Toso *et al*. [47].

**Figure F.3:**
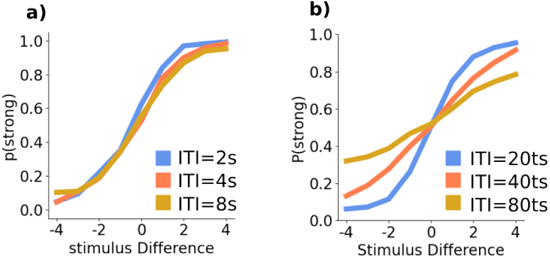
Time steps in our model: Example from the One Back paradigm. **a)** Conditioned psychometric curves based on the time elapsed between two stimuli (independent of the cloud). **b)** Same as **a)**, but for the model results.

#### F.2 The RNN model qualitatively reproduces behavioural effects of varying ISI

Experimental evidence shows that contraction bias depends on the time between stimuli – the longer the interval, the stronger the contraction bias. For instance, Akrami *et al*. [3] found that rats presented with pairs of auditory stimuli at varying ISIs (2, 4, 6 seconds) showed a characteristic time dependence in their psychometric curves: longer ISIs lead to less steep curves, as was reported in Fig. 1e of their paper. This behaviour is captured in our model, as shown in fig. C.1c.

We also found that our model correctly reproduces the behavioural effect of varying the inter-trial interval (ITI) in the One-Back paradigm with tactile stimuli, where the ITI is equal to the inter-stimulus interval. Despite differences in species and sensory modality, conditioned psychometric curves show the typical contraction bias effect, as shown in fig. F.3a), as the ITI is varied from 2 to 4 to 8. Our model, using proportionally varying ITIs (20, 40, 80 time steps), replicates this behaviour, as shown in fig. F.3b).

#### F.3 Neural vs. memory time scales

In our model, threshold linear units are a simplified representation of neuronal responses and are not directly mapped to intrinsic neuronal properties. They can represent individual neurons or the ensemble rate of a group of interconnected neurons. If we interpret the threshold linear units as modeling individual neurons, and considering the relationship between *γ* and synaptic conductances, then the time steps used in our simulations (e.g., 335 ms) would imply unrealistically large synaptic conductances on the order of seconds. One way to address this would be to either increase *γ* (which destabilises the simple neural model) or increase the number of time steps so that each corresponds to a duration between 1–4 ms, leading to a more realistic time scale of synaptic conductance. This would require a retuning of the model’s hyperparameters, for which a better theoretical understanding of the different regimes of our model would be of great help. However, this is out of the scope of the present work.

Here, we show that even with the present parameters, our model addresses the key challenge of modelling working memory, which is overcoming the disparity between neural time scales and memory duration. In our model, the duration for which a stimulus is held in memory and the rate at which its memory trace diverges depend not only on the neuronal time scale but also on the attractor’s shape and its ongoing modification through Hebbian learning solving the highlighted challenge. In fig. F.4a, we illustrate how the memory trace of different stimuli *S*_*A*_ evolves from their offset to the step just before the onset of *S*_*B*_, across all trials with the longest ISI (24 time steps) for a single model realization. The variation in the internal representation of the stimulus across trials as well as how fast they diverge results from both the activity state prior to stimulation and the strength of the recurrent connections. The vertical line at 10 time steps indicates the critical value that would be expected if memory were solely dependent on the neuronal time scale (assuming *γ* = 0.1), but no significant change occurs at this point. In fig. F.4b, where we average across trials and realizations, this effect becomes even more apparent. The memory of the stimulus persists well beyond the neuronal critical value, as the curves do not collapse either at 10 time steps or at 24.

**Figure F.4:**
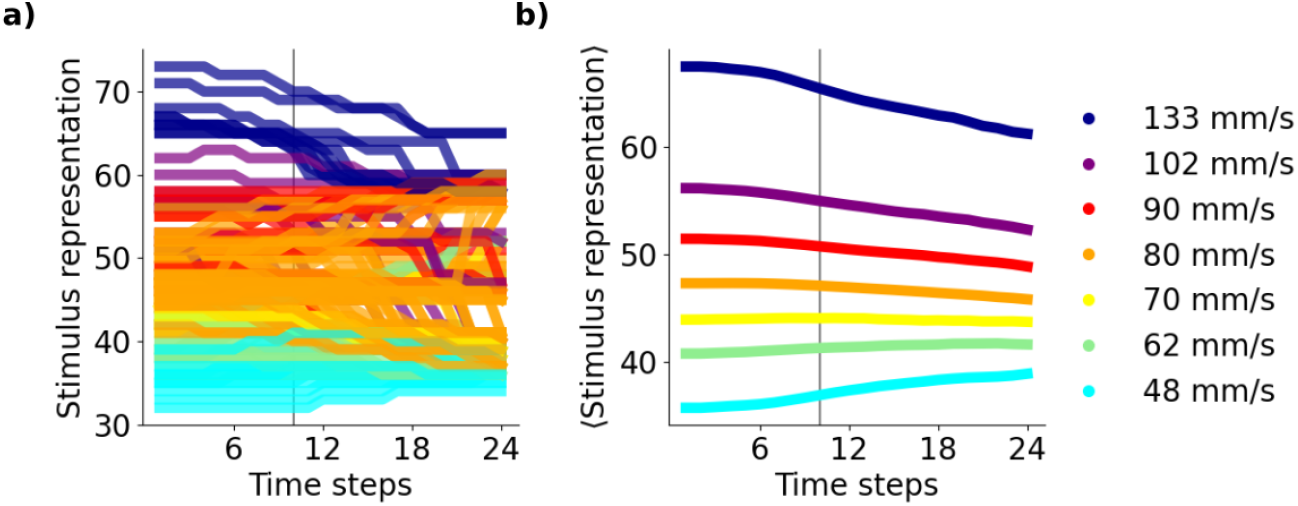
The effective timescale of memory. In a working memory paradigm with a model realization (using parameters from fig. F.1, second row, where ISI and stimulus duration align with experiments), we analyse trials with a 24-time step ISI. a) Shows individual trials, where memory decay varies due to prior stimuli and the neural state before *S*_*A*_. b) Displays the average across trials and realizations. The vertical line at 10 time steps indicates neuronal decay, yet memory persists beyond this point due to the attractor dynamics.

While neuronal time scales smooth the transition of activity and help maintain a stimulus representation, the free evolution of the system, in the absence of stimulation, is governed by the attractor, or the connections, which store the history of previous inputs. The basin of attraction can take different shapes—such as the steeper or smoother parabolas one could draw in fig. 3 e) and f) of the main text—which continuously change depending on the inputs and the system’s dynamics. A steeper basin causes the memory to decay more quickly, while a smoother one slows this process. Additionally, depending on the stimulus history, the representation of a stimulus may or may not align with the attractor in a given trial, leading to different memory divergence behaviours, despite the same neuronal time constant. Therefore, the memory time scale cannot be directly mapped to the neuronal time scale; it is instead an **effective time scale** that emerges from these factors and changes throughout the experiment.

